# Structural polymorphisms within a common powdery mildew effector scaffold as a driver of co-evolution with cereal immune receptors

**DOI:** 10.1101/2023.05.05.539654

**Authors:** Yu Cao, Florian Kümmel, Elke Logemann, Jan M. Gebauer, Aaron W. Lawson, Dongli Yu, Matthias Uthoff, Beat Keller, Jan Jirschitzka, Ulrich Baumann, Kenichi Tsuda, Jijie Chai, Paul Schulze-Lefert

**Affiliations:** Department of Plant Microbe Interactions, Max Planck Institute for Plant Breeding Research, 50829 Cologne, Germany; Institute of Biochemistry, University of Cologne, 50674 Cologne, Germany; Department of Plant and Microbial Biology, University of Zurich, 8008 Zurich, Switzerland; State Key Laboratory of Agricultural Microbiology, Hubei Hongshan Laboratory, Hubei Key Lab of Plant Pathology, College of Plant Science and Technology, Huazhong Agricultural University, 430070, Wuhan, China; Beijing Frontier Research Center for Biological Structure, Center for Plant Biology, School of Life Sciences, Tsinghua University, 100084 Beijing, China; Cluster of Excellence on Plant Sciences, Max Planck Institute for Plant Breeding Research, 50829 Cologne, Germany

**Keywords:** innate immunity, effector, crystal structure, powdery mildew, immune receptor

## Abstract

In plants, host–pathogen coevolution often manifests in reciprocal, adaptive genetic changes through variations in host nucleotide-binding leucine-rich repeat immune receptors (NLR) and virulence-promoting pathogen effectors. In grass powdery mildew (PM) fungi, an extreme expansion of a RNase-like effector family, termed RALPH, dominates the effector repertoire, with some members recognized as avirulence (AVR) effectors by cereal NLR receptors. We report the structures of the sequence-unrelated barley PM effectors AVR_A6_, AVR_A7_ and allelic AVR_A10_/AVR_A22_ variants, which are detected by highly sequence-related barley NLRs MLA6, MLA7, MLA10, and MLA22, and of wheat PM AVR_PM2_ detected by the unrelated wheat NLR PM2. The AVR effectors adopt a common scaffold, which is shared with the ribonuclease (RNase) T1/F1-family. We found striking variations in the number, position, and length of individual structural elements between RALPH AVRs, which is associated with a differentiation of RALPH effector subfamilies. We show that all RALPH AVRs tested have lost nuclease and synthetase activities of the RNase T1/F1- family and lack significant binding to RNA, implying that their virulence activities are associated with neo-functionalization events. Structure-guided mutagenesis identified six AVR_A6_ residues that are sufficient to turn a sequence-diverged member of the same RALPH subfamily into an effector specifically detected by MLA6. Similar structure-guided information for AVR_A10_ and AVR_A22_ indicates that MLA receptors detect largely distinct effector surface patches. Thus, coupling of sequence and structural polymorphisms within the RALPH scaffold of PMs facilitated escape from NLR recognition and potential acquisition of diverse virulence functions.

## Introduction

Interactions of plants with host-adapted pathogens are often subject to a population-level arms race involving competing sets of co-evolving genes encoding plant immune receptors and pathogen effectors, the former being essential components for plant immunity and the latter being required for pathogen virulence (1). Intracellular nucleotide-binding domain leucine-rich repeat-containing receptors (NLRs) in plants are key components of the plant immune system and typically detect strain-specific pathogen effectors, known as avirulence (AVR) effectors, to activate immune responses that terminate pathogen proliferation. Canonical plant NLRs share a tripartite domain organization consisting of a variable N-terminal signaling domain, a central nucleotide-binding oligomerization (NOD) domain, followed by a C-terminal leucine-rich repeat region (LRR) with or without a Jelly roll/Ig-like (JID) domain (2–5). The majority of these NLRs carry either a coiled-coil (CC) domain or a Toll-interleukin 1 receptor (TIR) domain at the N-terminus and are termed CNLs or TNLs, respectively (2, 3). Pathogen effector perception by NLRs can occur via diverse mechanisms, including direct effector binding to polymorphic LRR and C-JID domains (4–7). Pathogen recognition can also be indirect, with NLRs detecting pathogen effector-mediated modifications of host proteins (guardees) or mimics of these proteins (decoys), including decoys integrated into NLRs (8, 9). Upon effector-mediated activation, canonical CNLs and TNLs undergo extensive structural inter-domain rearrangements and oligomerization to form resistosomes composed of five or four NLR protomers, respectively. CNL resistosomes integrate into host cell membranes to act as calcium-permeable channels, whereas TNL resistosomes produce nucleotide-based second messengers for immune signaling (4–7, 10, 11). Ultimately, these NLR-mediated immune responses often result in regulated local death of host cells at sites of attempted pathogen ingress, the so-called hypersensitive response.

In host-adapted pathogens, co-evolution with their hosts occurs in iterative cycles and has resulted in genomic expansion of the plant NLR arsenal as well as the pathogen effector complement (1). *NLR* genes in plants are often organized in complex clusters of paralogous genes, and several examples of allelic series of *NLR*s have been reported in host populations, with each receptor variant conferring a different effector recognition specificity. The repertoire of effector genes of pathogenic fungi is much larger (typically hundreds) compared to pathogenic bacteria (a few dozen) and the effectors are often lineage- or species-specific innovations, suggesting that effectors of different fungal lineages evolve rapidly and independently of each other (12, 13). Sequence-relatedness between individual fungal effector genes is often low or undetectable. However, there is increasing evidence that many of these effectors are structurally related. Thus, it is possible that the effector repertoire of pathogenic ascomycetes consists of a limited number of structural folds (12, 14–21). Yet, it is still unclear whether each effector fold is associated with a common biochemical function or serves as a scaffold for diversified virulence activities.

The powdery mildews *Blumeria gramini*s f. sp. *hordei* (*Bgh*) and *Blumeria graminis* f. sp. *tritici* (*Bgt*) infect monocotyledonous barley and wheat, respectively, and are widespread, obligate biotrophic ascomycete fungi. *Bgh* and *Bgt* each secrete hundreds of candidate effector proteins (CSEPs) to promote pathogen growth. Numerous allelic CNL variants are encoded at the barley *Mla* or wheat *Pm2* or *Pm3* resistance loci, each conferring isolate-specific immunity to *Bgh* or *Bgt* strains, respectively, with matching AVR effectors (22–29). Although MLA, PM2 and PM3 are phylogenetically unrelated CNLs, receptors encoded by each of these loci with different resistance specificities share >90% sequence identity. Together this indicates that these polymorphic CNLs in barley and wheat contribute to co-evolution with *Bgh* and *Bgt*, respectively. The sequence-unrelated paralogous *Bgh* avirulence effectors AVR_A1_, AVR_A6_, AVR_A7_, AVR_A9_, AVR_A13_ and the sequence-related allelic variants AVR_A10_/AVR_A22_ are likely to be recognized directly by barley MLA1, MLA6, MLA7, MLA9, MLA13, MLA10 and MLA22, respectively, through their polymorphic LRR domains (22, 25, 29). AVR_PM2_, AVRPM3^A2/F2^, AVRPM3^B2/C2^, and AVRPM^3D^ were identified in *Bgt* and were shown to be recognized by wheat PM2a and PM3a/PM3f, PM3b/PM3c, and PM3d, respectively, with PM2a and PM3 LRRs also functioning as recognition specificity determinants (27, 28, 30).

Genome-wide AlphaFold2 (AF2) modeling of fungal effector complements identified extreme expansion of lineage-specific, sequence-unrelated, structurally similar effector families in *B. graminis* and the rust fungus *Puccinia graminis* (20). This modeling predicted that at least 70% of all *Bgh* effectors adopt the common fold of RNase-like proteins associated with haustoria (RALPHs) (22, 23, 25–31). RALPH effectors in a given *Bgh* or *Bgt* strain are typically encoded by >400 paralogous genes organized in at least 15 RALPH subfamilies, with no detectable sequence similarity between subfamilies (20, 32–35). All 14 identified *AVR* effectors in *Bgh* and *Bgt* encode variants of predicted RALPH effectors. The structure of a CSEP with unknown avirulence activity, *Bgh* CSEP0064, features an RNase-like fold and is thought to act as a pseudoenzyme that binds to host ribosomes, thereby inhibiting the action of toxic plant ribosome-inactivating proteins (31). Other modelled RALPH effectors with unknown avirulence activity interact with different barley proteins *in vitro* and *in vivo* (36–38). *Bgh* AVR_A1_ and the predicted RALPH effector CSEP0491 interact with the barley endoplasmic reticulum-localized J-domain-containing protein *Hv*ERdj3 (39).

We report here the crystal structures of four *Bgh* AVR_A_ effectors and of *Bgt* AVR_PM2_ after heterologous expression and purification from *E. coli* or insect cells. All five AVR effectors adopt the common RALPH scaffold, but they have striking structural differences associated with differentiation of RALPH effector subfamilies. Using biochemical assays, we confirm that all AVR RALPHs tested have lost catalytic activities of the ribonuclease T1/F1-family. The AVR_A6_ structural template was used for mutagenesis of a RALPH effector belonging to the same subfamily to construct MLA6 gain-of-recognition hybrid effectors upon expression in barley protoplasts and heterologous *N. benthamiana*. Six amino acid substitutions were sufficient to turn the sequence-diverged effector CSEP0333 into a variant specifically recognized by MLA6. Our findings suggest that coupling of sequence and structural polymorphisms within the RALPH scaffold facilitated both escape from CNL receptor recognition and potential acquisition of new virulence functions, which might explain the proliferation and overabundance of this effector family in *B. graminis*.

## Results

### *Blumeria graminis* AVR effectors adopt a common structural scaffold

To better understand the co-evolution of AVR effectors of *Bgh* with matching barley MLA receptors, we sought to obtain three-dimensional effector structures using X-ray crystallography. To extend this analysis to AVR effectors belonging to the RALPH effector superfamily in a reproductively isolated *B. graminis* lineage (30), we included the wheat powdery mildew effector AVR_PM2_, which is detected by wheat Pm2a (24, 30). All AVR effectors were recombinantly expressed without their predicted signal peptides and throughout this manuscript we refer to the residue positions in the mature proteins starting with methionine. We obtained well-diffracting crystals for AVR_A6_, AVR_A7_, AVR_A10_, AVR_A22_ of *Bgh* and AVR_PM2_ of *Bgt* and solved their structures with molecular replacement (**Fig. S1**). The data processing and refinement statistics for the structures are outlined in **Table S1**.

The cores of the five *B. graminis* AVR effector proteins are composed of two β-sheets and a central α-helix (**Fig. 1A**). The first ß-sheet is formed by two or three anti-parallel ß-strands, of which two contribute an N-terminal ß-hairpin (AVR_A7_, AVR_A10_, AVR_A22_, AVR_PM2_), whereas another ß-strand is at the very C-terminus (AVR_A6_, AVR_A10_, AVR_A22_ and AVR_PM2_). The second ß-sheet is formed by three or four anti-parallel ß-strands, of which at least two pack against the α-helix to stabilize the conformation of the ß-sheet (**Fig. 1A**). The long loop region following the α-helix is stabilized by polar contacts with the central second ß-sheet. Two conserved cysteine residues form an intramolecular disulfide bridge that connects the N- and C-terminal ends of the AVR proteins (**Fig. 1B**).

**Fig. 1.**
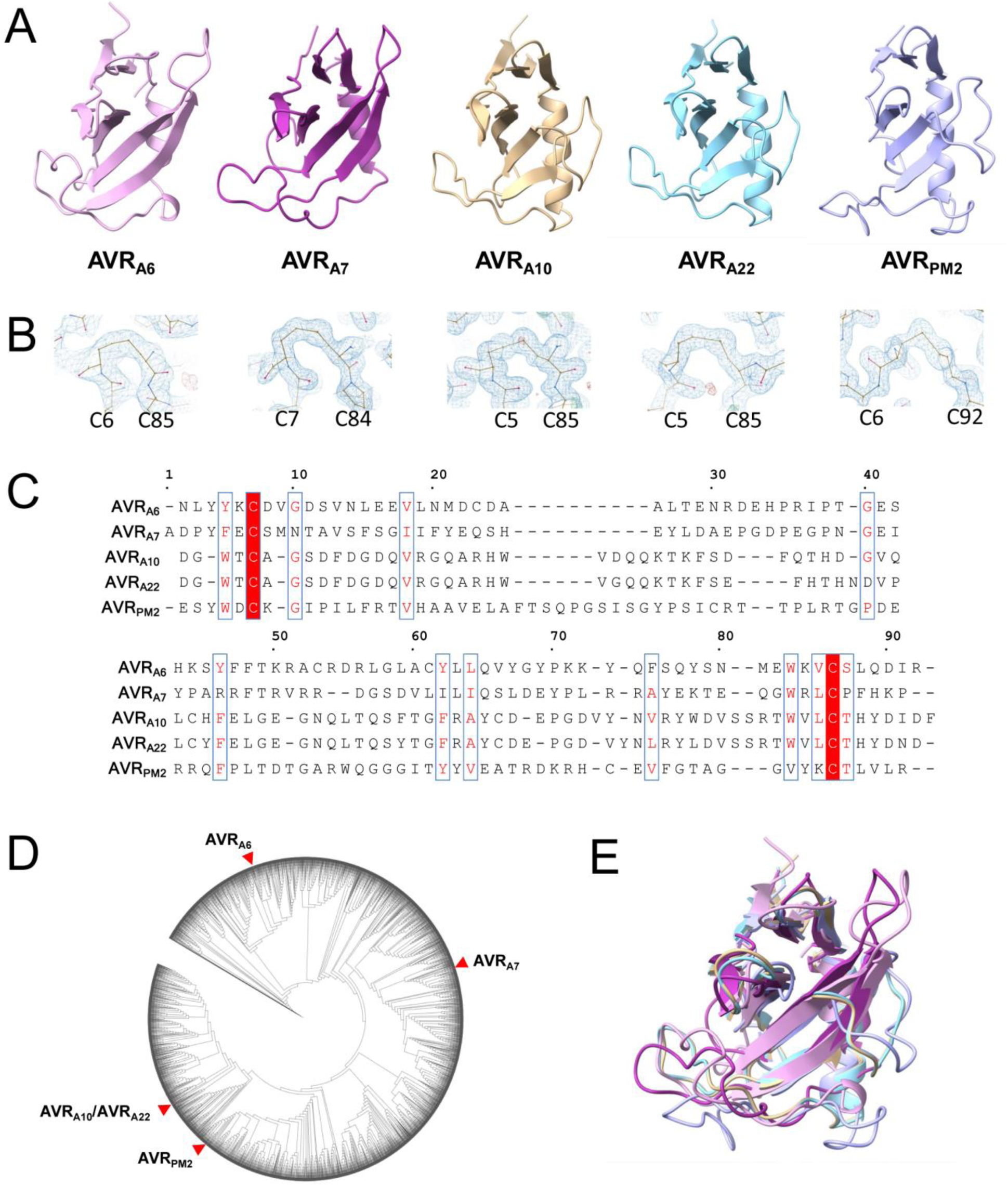
*Blumeria graminis* AVR effectors adopt a common structural scaffold. (A) Cartoon representation of the crystal structures of AVR_A6_, AVR_A7_, AVR_A10_, AVR_A22_ and AVR_PM2_. The effectors exhibit a canonical (α+β) RNase-like fold. (B) Disulfide bonds are conserved in *Blumeria* AVRs. AVR_A6_, AVR_A7_, AVR_A10_, AVR_A22_ and AVR_PM2_ form intramolecular disulfide bridges that connect the N and C termini. The disulfide bridge is indicated in the density map. (C) Amino acid sequences alignment of AVR_A6_, AVR_A7_, AVR_A10_, AVR_A22_ and AVR_PM2_ without signal peptides. Red background indicates amino acid similarity. The alignment was generated using ESPript 3.0 (60). (D) Maximum likelihood phylogeny including all predicted CSEPs from *B. graminis* f. sp. *poae, lolium, avenae, tritici* 96224*, hordei* DH14*, secalis* S1459*, triticale* T1-20, and *dactylidis*. AVR_A6_, AVR_A7_, AVR_A10_, AVR_A22_ and AVR_PM2_ are widely separated in the phylogeny. (E) Superposition of AVR_A6_, AVR_A7_, AVR_A10_, AVR_A22_ and AVR_PM2._

Except for allelic AVR_A10_ and AVR_A22_, the sequence similarity between the five AVR effectors is extremely low (maximally 40% similarity and 19% identity; **Fig. 1C**). Furthermore, based on multiple sequence alignment of the available *B. graminis* effector complement, they represent distinct effector subfamilies and are widely separated on the corresponding maximum likelihood phylogenetic tree (**Fig. 1D**). We searched the DALI database of known structures and found that all AVR effectors adopt the common RALPH scaffold (**Fig. 1E)** (20). Sequence conservation among the AVR effectors is limited to a few residues that are hydrophobic and buried in the cores of the structures. Previous studies have identified the F/Y/WxC motif as a common feature of powdery mildew effectors, based on sequence similarity analysis (40, 41). The aromatic residue of the F/Y/WxC motif is buried in the core and forms van-der-Waals contacts with residues in ß5 in AVR_A6_ and AVR_A7_ or ß6 in AVR_A10_/AVR_A22_ and AVR_PM2_. Therefore, the F/Y/WxC motif contributes to stabilizing the common RALPH fold. A similar role can be assigned to other conserved hydrophobic residues including V18, Y45, Y61, L63, W82 and V84 of AVR_A6_ (**Fig. 1C**), suggesting that despite the overall extreme sequence divergence of the RALPH effector family, evolutionary selection has also favored the conservation of less than 20% of residues (**Fig. 1C**), largely scattered in the primary sequence, to maintain a stable common structural scaffold.

### Structural variations of *B. graminis* AVR effectors

A structure-based similarity search was carried out using the DALI server (42). As anticipated, AVR_A6_, AVR_A7_, AVR_A10_, AVR_A22_ and AVR_PM2_ are structurally related to the ribonuclease T1/F1 family, despite the lack of detectable relatedness in their protein sequences (**Fig. S3**). The overall structures of AVR_A10_, AVR_A22_ and AVR_PM2_ are more similar to RNase T1 than to AVR_A6_ and AVR_A7_, indicating that structural variation among RALPH AVR effectors within a reproductive lineage of *B. graminis* exceeds the variation relative to the ribonuclease T1/F1 family. Similarly, AVR_PM2_ in *Bgt* is structurally more similar to AVR_A10_ and AVR_A22_ in *Bgh* than to AVR_A6_ and AVR_A7_ in *Bgh*, showing that structural dissimilarity between RALPH AVR effectors within a reproductive lineage of *B. graminis* can be greater than between two powdery mildew lineages (**Fig. 2B**). Interestingly, the structural relationship among the AVR effectors is consistent with their phylogenetic relationship. Namely, AVR_A10_/AVR_A22_ and AVR_PM2_ are located distant to AVR_A6_ and AVR_A7_ in the phylogenetic tree of all *Blumeria* CSEPs (**Fig. 1D**).

**Fig. 2.**
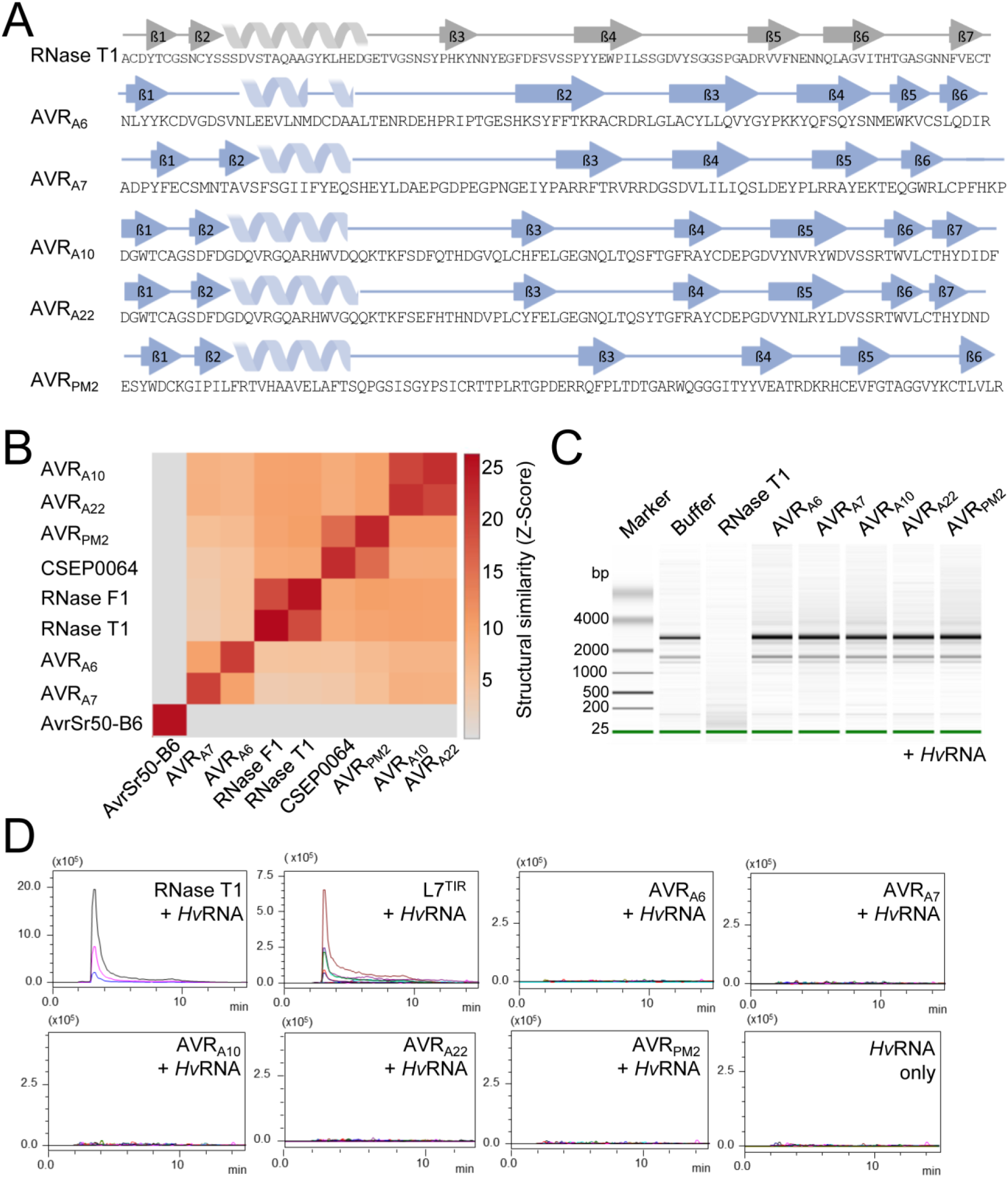
*Blumeria graminis* AVR effectors are pseudo-RNases with diversified structural features. (A) *B. graminis* AVR and RNase T1 (9RNT) proteins harboring diversified secondary structural features. ß-strands are indicated by arrows, α-helices by spirals. Secondary structures are pictured according to ChimeraX using BioRender. (B) Pairwise comparison between crystal structures using Dali server (42). (C) Recombinant AVR effector proteins lack ribonuclease activity. AVR effectors (1 µM) were co-incubated with *Hv*RNA and then analyzed on a Bioanalyzer to evaluate RNA degradation. (D) RNase-like AVR effectors lack 2’3’-cNMP synthetase activity. Samples were subjected to LC-MS/MS for metabolite identification and quantification.

Although the AVR effector proteins share a general structural similarity with RNase T1, there are striking local structural variations between these proteins. For example, compared with RNase T1, all AVR effectors except AVR_A7_ possess an additional ß-strand (ß6 in AVR_A6_ and AVR_PM2_, ß7 in AVR_A10_/AVR_A22_) after the disulfide bridge-forming cysteine in the C-terminus (**Fig. 2A; Fig. S5**). Variations in individual structural elements are also evident between the AVR RALPH effectors. The first ß-sheet of AVR_A10_/AVR_A22_ and AVR_PM2_ consists of three ß-strands, whereas the corresponding ß-sheet in AVR_A6_ and AVR_A7_ consists of only two ß-strands. In AVR_A7_, the two ß-strands of the first ß-sheet are located in the N-terminus, whereas in AVR_A6_ both N- and C-termini contribute one ß-strand each. In AVR_A6_ and AVR_A7_, the ß-strands of the central ß-sheet that face the α-helix are substantially longer than the corresponding ß-strands in AVR_A10_, AVR_A22_ and AVR_PM2_. In addition, the loop region connecting ß3 and ß4 of AVR_A7_ is much shorter than its equivalent in AVR_A6_. Finally, the length of the α-helix is also variable among the AVR effectors, except for allelic AVR_A10_ and AVR_A22_ (**Fig. 1A; Fig. 2A**). Collectively, this indicates an unexpected structural diversification among AVR RALPH effectors, which is associated with a differentiation of RALPH effector subfamilies despite the maintenance of a common structural scaffold.

Superimposition of RNase T1 (PDB: 9RNT) with AVR_A6_, AVR_A7_, AVR_A10_, AVR_A22_ and AVR_PM2_ illustrates structural homology, but the predicted residues for RNA hydrolysis are not conserved in the AVR effectors (**Fig. S3**). We confirmed previous studies that suggested (29, 31, 41) that AVR effectors are pseudo-RNases that cannot hydrolyze total barley RNA (*Hv*RNA) (**Fig. 2C; Fig. S4**). RNase T1 has also the capacity to produce 2’, 3’-cyclic nucleotide monophosphates (mainly 2’, 3’ -cGMP), which are putative second messengers in TNL-mediated mediated plant immunity (43). To test whether the *B. graminis* AVR effectors are catalytically active in producing 2’, 3’ -cNMP, we co-incubated the RNase T1, L7^TIR^, and powdery mildew AVR effectors with *Hv*RNA as substrate. Liquid chromatography coupled with mass spectrometry (LC-MS) analysis showed that only RNAse T1 and L7^TIR^ could synthesize 2’,3’-cAMP/cGMP under these conditions (**Fig. 2D**). To test whether the RNase-like effector proteins can bind *Hv*RNA, we performed microscale thermophoresis (MST) experiments. However, the affinity of the AVR effectors for total RNA was not significantly different to the measured affinity of non-RNase-like fold proteins (BSA, GST and AvrSr50 from *P. graminis* f. sp. *tritici* (**Fig. S4C**). This strongly suggests that RALPH AVR effectors have lost known enzymatic activities of the RNase T1/F1 family and exhibit variation in the number, position, and length of individual structural elements. This raises the possibility that upon escape from CNL detection, their potential virulence functions were associated with neo-functionalization events involving combined sequence and structural diversification within the common RALPH scaffold.

### Structural determinants of AVR_A6_ detection by MLA6

CSEP0333 is a member of the AVR_A6_ effector family (family E008) but is not recognized by MLA6 (29, 35). Based on their significant sequence identity (60%), we suspected that this effector had a similar structure to AVR_A6_ and reasoned that substituting residues in CSEP0333 with those of AVR_A6_ may convert this family member into an effector recognized by MLA6. Based on the crystal structure, AVR_A6_ was divided into three units: an N-terminal part comprising the ß1 and the α-helix (residues 1–26), a central segment that includes the long loop region and ß2 (residues 27–53), and a C-terminal part that includes the ß-strands ß3–ß6 (residues 54–91). Each of these segments was swapped with the corresponding unit in AVR_A6_ or CSEP0333, resulting in six AVR_A6_/CSEP0333 effector chimeras (**Fig. 3A**). We then individually co-expressed the hybrid effectors together with MLA6 in barley protoplasts and quantified cell death by measuring luciferase reporter activity (44). Swapping of either the N-terminal or C-terminal segment of AVR_A6_ with the corresponding unit in CSEP0333 (constructs A6N and A6C, respectively) did not lead to loss of recognition by MLA6 (**Fig. 3C**). As anticipated, CSEP0333 with its N-terminal or C-terminal part substituted with the equivalent parts of AVR_A6_ (constructs B6N and B6C, respectively) did not trigger MLA6- dependent cell death. However, when the central segment was exchanged to that in CSEP0333 (A6M), cell death was completely abolished. Strikingly, swapping the central segment of CSEP0333 with that of AVR_A6_ was sufficient to activate MLA6-mediated cell death in barley protoplasts (**Fig. 3A**), i.e., there was a gain in effector recognition. *Agrobacterium*-mediated individual co-expression of the six effector hybrids with a C-terminal mYFP-tag together with MLA6 in heterologous *N. benthamiana* produced comparable differential cell death phenotypes (**Fig. 3B**). All effector proteins were detectable in *N. benthamiana* leaf extracts (**Fig. 3C**). This demonstrates that effector detection by the receptor does not require a second barley protein.

**Fig. 3.**
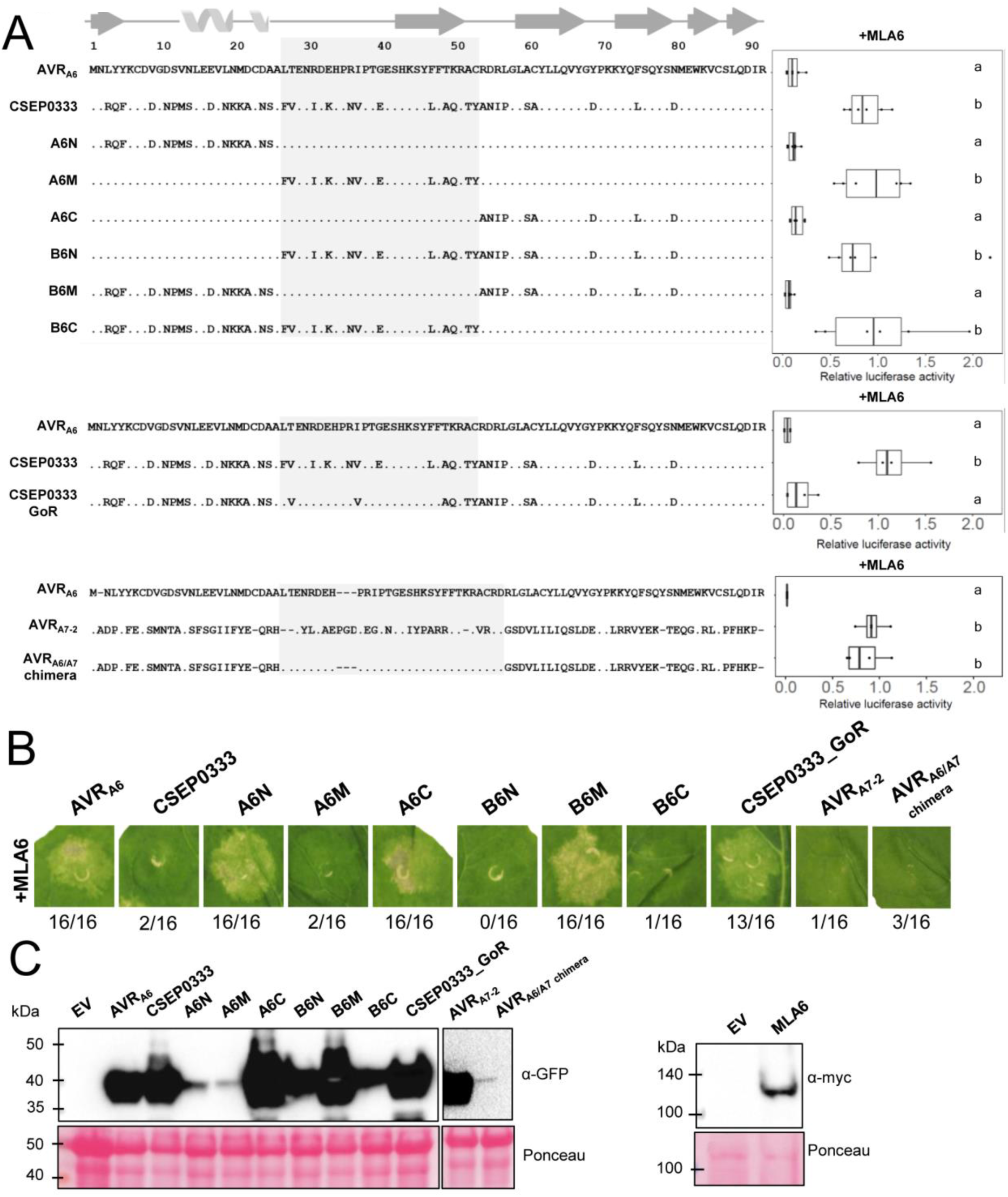
Six amino acids in the central segment of AVR_A6_ are essential for detection by MLA6. (A) Chimeric effectors were co-expressed with MLA6 in barley protoplasts and cell death was quantified by measuring luciferase reporter activity. Letters indicate results of statistical variance analysis using Kruskal-Wallis test followed by Dunn’s post hoc tests (P < 0.05). Raw relative luciferase measurements and P-values for all protoplast plots are provided in Supplementary Data S6. (B) *Agrobacterium*-mediated co-expression of the six effector chimeras with a C-terminal mYFP-tag together with MLA6 in *N. benthamiana* produced comparable differential cell death phenotypes. (C) All effector proteins were detectable in *N. benthamiana* leaf extracts, except for spurious amounts of the AVR_A6/A7_ chimera.

To define which part of the 12 residues in the central segment of AVR_A6_ are important for recognition by MLA6, we subdivided the central segment into two parts that encompass the loop region (seven polymorphic residues) or the ß2 and loop region connecting ß2–ß3 (five polymorphic residues). We found that AVR_A6_ hybrids in which either of these two parts are swapped with the respective unit present in CSEP0333 (constructs M1 and M2) are not recognized by MLA6 (**Fig. S6 & S7**). This indicated that residues from both the loop regions as well as ß2 are involved in AVR_A6_ recognition by MLA6. Therefore, we used the constructs M1 and M2 as templates for site-directed mutagenesis to create 19 additional AVR_A6_ higher-order mutant constructs with different combinations of targeted amino acid substitutions (**Fig. S6 & S7**). By comparing cell death induced by different combinations of amino acid substitutions, it was possible to identify residues that are essential for recognition by MLA6. For example, the construct M1^F27L/K33E/N36R/E40G^ did not lead to induction of cell death even though the protein was detectable in *N. benthamiana* leaf extracts, whereas the construct M1^F27L/I31R/K33E/N36R/E40G^ was able to reduce LUC activity to a level comparable with wild-type AVR_A6_ (**Fig. S6 & S7**). These results indicate that L31R can rescue the phenotype generated by the other four mutations. Cell death induction after co-transfection with MLA6 in protoplasts was only observed for constructs that include the six residues L27, R31, E33, R36, G40 and F47 of AVR_A6_ (**Fig. S6 & S7**). We then introduced the six amino acid substitutions F27L, I31R, K33E, N36R, E40G and L47F in CSEP0333, and the resulting construct, termed CSEP0333^GoR^ (CSEP0333^Gain-of-Recognition^), was able to trigger MLA6- dependent cell death both in barley protoplasts and *N. benthamiana* (**Fig. 3A, 3B**). All constructs that induce cell death when co-expressed with MLA6 were also co-expressed together with MLA7 in barley protoplasts. Only AVR_A7_ induced MLA7-dependent cell death, confirming recognition specificity of the tested hybrid effectors by MLA6 (**Fig. S6 & S7**). In summary, our results show that six amino acid substitutions in the central segment of CSEP0333 are sufficient to turn this sequence-diverged effector into a variant specifically recognized by MLA6. It is therefore possible that CSEP0333 evolved from AVR_A6_ by immune evasion.

Among the five resolved structures of AVR effectors, AVR_A6_ is structurally most similar to AVR_A7_ (**Fig. 2B**). Nevertheless, based on sequence relatedness, AVR_A7_ belongs to effector family 29, whereas AVR_A6_ and CSEP0333 both belong to effector family 8 (35). We constructed a chimeric effector in which the central segment of AVR_A7_ is swapped for the equivalent part of AVR_A6_ (AVR_A6/A7_ chimera). However, co-expression of the AVR_A7_/AVR_A6_ hybrid with MLA6 did not induce cell death either in barley protoplasts or *N. benthamiana*. The AVR_A7_/AVR_A6_ chimera also did not cause MLA7-dependent cell death in barley protoplasts, and the hybrid protein was barely detectable in *N. benthamiana* leaf extracts (**Fig. 3; Fig. S6 & S7**). This shows that the RALPH interfamily AVR_A7_/AVR_A6_ hybrid is unstable *in planta*, presumably due to the structural differences between wild-type AVR_A6_ and AVR_A7_, which likely hinder the proper folding of the hybrid protein.

### MLA6, MLA10, MLA22 and PM2 CNLs each recognize largely different surface patches on the RALPH scaffold

Previously, four amino acids were identified in AVR_A10_ that cannot be exchanged to the respective residues in AVR_A22_ without abrogating recognition by MLA10 and five amino acids in AVR_A22_ that cannot be exchanged to the corresponding residues in AVR_A10_ without losing MLA22-dependent recognition (29). Furthermore, amino acids that constitute the ‘head epitope’ of AVR_PM2_ are important for specific recognition by wheat PM2a (24). When mapped onto the structures of these effector proteins, these epitopes are located at different sites (**Fig. 4**). In AVR_A10_, one residue is located in the loop region after the α-helix (D33), one residue is in the loop region between ß3 and ß4 (F57) and two map to the ß-strands ß3 and ß5 (H44 and W76, respectively). By contrast, in AVR_A22_ the residues important for the recognition by MLA22 are mainly located in the loop region between ß2 and ß3 (H35, N38, D39 and P41). The residue G25 is located at the end of the α-helix. In AVR_A6_, the six amino acids that are essential for recognition by MLA6 are located in a surface patch at the loop region between the α-helix and ß3. All identified residues are highly surface-exposed (**Fig. S8**). Given that MLA receptors likely detect cognate AVR RALPH effectors directly (22), these findings indicate that the interface between an AVR_A_ effector and an MLA receptor is different for each of these matching effector–receptor pairs (see below).

**Fig. 4.**
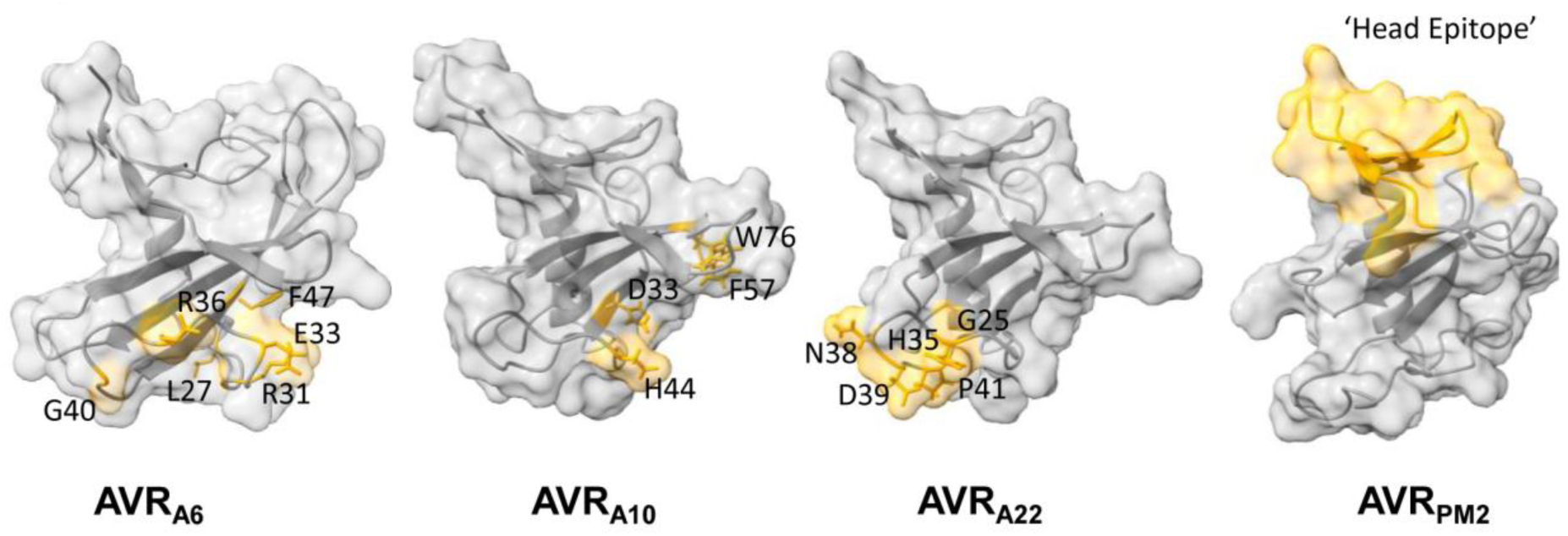
MLA6, MLA10, MLA22 and PM2 CNLs each recognize largely distinct surface patches on the common RALPH effector scaffold. Locations of residues in AVR_A6_, AVR_A10_, AVR_A22_ and AVR_PM2_ that determine the respective MLA6, MLA10, MLA22 and PM2a recognition specificities are highlighted in orange color. The residues of AVR_A10_ and AVR_A22_ required for specific MLA10 and MLA22 recognition as determined in (29). The residues of AVR_PM2_ important for recognition of AVR_PM2_ were determined in (24).

### RALPH effector subfamilies harbouring avirulence effectors have different conserved surface arrays

AVR_A6_, AVR_A7_, AVR_A10_/AVR_A22_ and AVR_PM2_ belong to four distinct phylogenetic effector subfamilies (**Fig. 1D**). We sought to investigate the evolution of these RALPH effectors in their respective subfamilies, by mapping highly polymorphic as well as conserved residues onto the resolved AVR structures. We then highlighted conserved residues (>70%) in the crystal structures (**Fig. 5**). Most conserved residues are buried in the core of the structure and contribute to maintain the structural scaffold. Similarly, most surface-exposed residues are highly polymorphic. Interestingly, however, some residues are conserved in the respective subfamily, although they have a high relative solvent-accessibility (SA). For example, in AVR_A6_ and AVR_A7_, in the loop region after the α-helix, three highly exposed residues are conserved: proline at position 38 in AVR_A6_ (64% SA; 30 out of 33 family members) or position 39 in AVR_A7_ (72% SA; 71 out of 72 family members); glycine at position 40 in AVR_A6_ (73% SA; 24 out of 33 family members) or position 41 in AVR_A7_ (46% SA; 67 out of 72 family members); glutamic acid at position 41 in AVR_A6_ (42% SA; 26 out of 33 family members) or position 42 in AVR_A7_ (37% SA; 44 out of 72 family members) (**Fig. 5A; Supplementary File 1**). This surface patch appears not to be conserved in AVR_A10_/AVR_A22_ and AVR_PM2_, in which the conserved surface-exposed residues are not confined to a discrete surface patch but rather map to multiple locations.

**Fig. 5.**
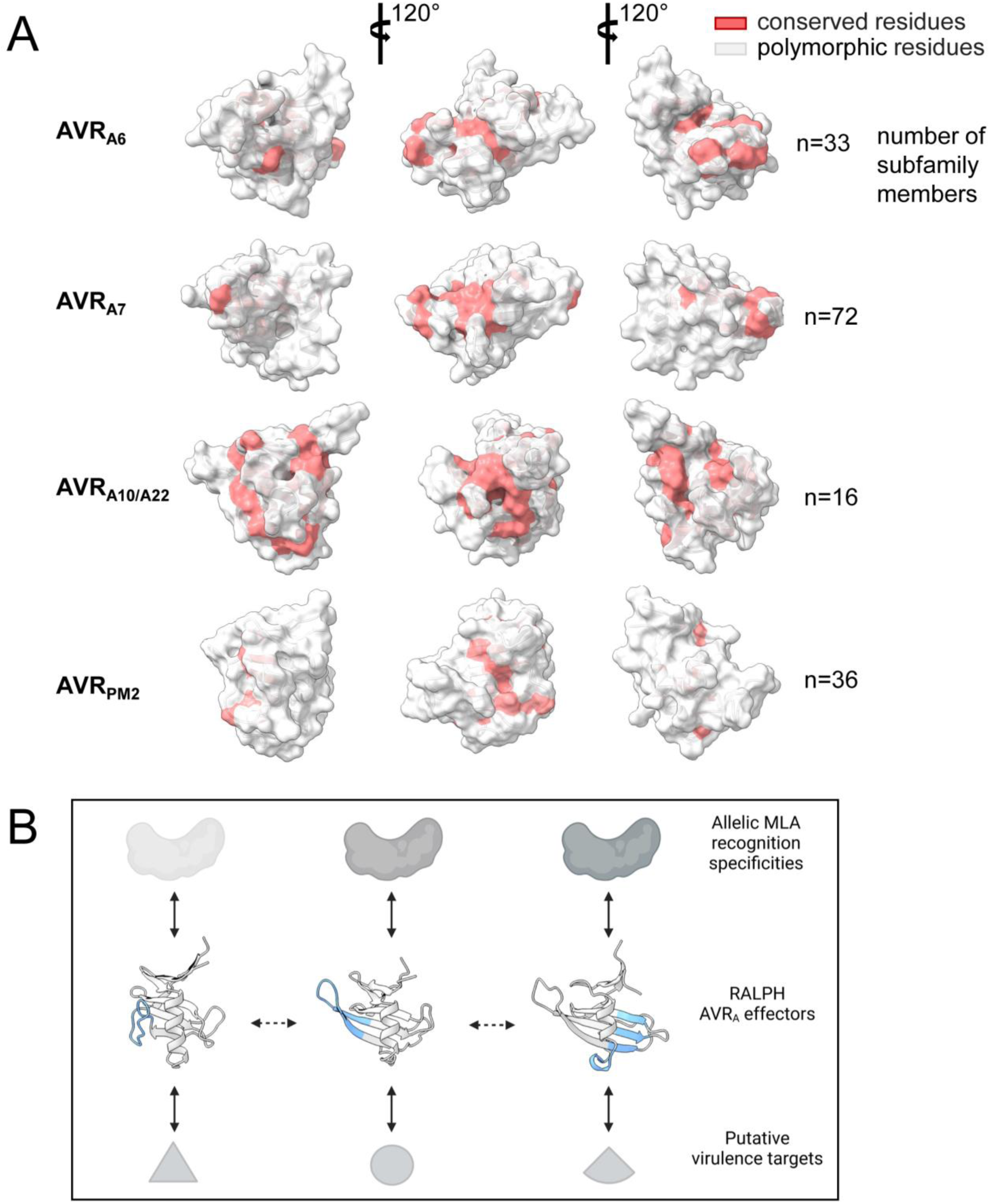
RALPH effector subfamilies harboring avirulence effectors have overlapping or distinct conserved surface arrays. (A) All CSEPs from *B. graminis* f sp *poae, lolium, avenae, tritici* 96224, *hordei* DH14, *secalis* S1459, *triticale* T1-20, and *dactylidis* were subjected to BLASTP. CSEPs that share >30% sequence identity and similar size to the crystallized RALPH AVR effectors were retained for further analysis using Muscle. Red color indicates conserved (70% threshold) residues. (B) Model for MLA receptor and RALPH effector co-evolution. Major local structural differences between RALPH AVR_A_ effectors are highlighted in blue. Solid bidirectional arrows indicate selection pressure by co-evolving protein pairs, dashed bidirectional arrows represent adaptive genetic changes in RALPH effectors.

## Discussion

We have resolved here five crystal structures of AVR effectors from a total of 14 RALPH AVR proteins of *B. graminis*, each of which was purified to homogeneity and in sufficient quantity (**Fig. S1 & Fig. S2**). It is possible that the extensive diversity of structural elements found among the crystallized RALPH AVR proteins, combined with the exceptional sequence diversification of surface-exposed residues, makes it challenging to obtain well-diffracting crystals for the remaining 9 AVR effectors (**Fig. S2**). This could explain why a wide range of different conditions had to be screened for successful crystallization of the purified proteins, although they share one common structural scaffold (Methods). Another factor could be that some RALPH AVR effectors adopt a stable conformation only in association with their effector targets inside host cells.

Recently, machine-learning algorithms for the prediction of protein structures, such as AF2, have greatly increased in accuracy. However, AF2 is homology-based, and the accuracy of the predictions depends on available experimental structures in the multiple sequence alignment (MSA) (45). The crystal structure of avrSr50-B6 of *P. graminis* f. sp. *tritici* revealed that the model predicted by AF2 is largely incorrect (46). This is consistent with the resolved powdery mildew AVR structures. While the topology of predicted AVR_A6_, AVR_A7_ and AVR_PM2_ is close to the real structures, the predictions of AVR_A10_ and AVR_A22_ are largely inaccurate. When the crystal structures of AVR_A10_/AVR_A22_ are compared to the predicted AF2 model, only 55 atoms (from a total 97 atoms) can be aligned, but with a root-mean-squared deviation (RMSD) of 4.50 Å and 4.90 Å, respectively (**Fig. S10**). AVR_PM2_ of *Bgt* is in the same family as the previously resolved CSEP0064 of *Bgh*, which can explain why a subset of effectors including AVR_PM2_ are predicted more accurately than AVR_A10_/AVR_A22_. AVR_A10_/AVR_A22_ belong to a relatively small effector subfamily (**Fig. 1D; Fig. S9**). Thus, our work on the structural diversification among RALPH AVR effectors illustrates that MSA-based modeling algorithms must be applied with greater caution, because the models captured only a subset of the actual diversity of individual structural elements within a common scaffold.

Despite mounting evidence that fungal effectors with low or undetectable sequence similarity may be structurally related and may share a common scaffold (12, 14–17, 20), it is still unclear whether this indicates shared scaffold-specific biochemical activity. We have provided evidence that all five RALPH AVR effectors tested have lost both RNase and synthetase catalytic activities of the evolutionarily ancient ribonuclease T1/F1 family. Furthermore, contrary to a previous report on the CSEP0064 RALPH effector from *Bgh* (31), none of the five AVR RALPH effectors tested in this study has an affinity for RNA that is significantly different to the measured affinity of non-RNase-like fold proteins, including the AvrSr50 effector from *P. graminis* f. sp. *tritici*. Our findings rather suggest that virulence activities of RALPH-AVR effectors belonging to different subfamilies are the result of neofunctionalization events, which could explain why several tested RALPH effectors were found to interact with different host proteins (31, 36–39). Evolutionary selection of the conserved RALPH framework may indicate existence of non-variable functions such as translocation across the fungal and host membranes into host cells and/or stable folding.

Sequence-based *Bgh* genome-wide analysis (35), as well as large-scale structural modeling using AF2 indicate that approximately 70% of candidate secreted effector proteins are RALPHs (20). We have been able to generate, by only six amino acid substitutions, a gain-of-recognition variant of the CSEP0333 effector specifically detected by MLA6 and belonging to the subfamily containing AVR_A6_, indicating that CSEP0333 may have evolved from AVR_A6_ by immune evasion. The AVR_A_ effectors AVR_A6_, AVR_A7_, AVR_A9_, AVR_A13_, and the allelic variants AVR_A10_/AVR_A22_ each belong to a different RALPH subfamily with multiple paralogs (22, 25, 29). This suggests that, with the exception of MLA10 and MLA22, each MLA recognition specificity drives sequence diversification within a different RALPH subfamily. Recent structural information on CNL and TNL resistosomes in *A. thaliana*, *N. benthamiana* and wheat, all activated by direct AVR interactions, has shown that multiple surface-exposed effector residues make extensive contacts with their respective neighboring residues in C-terminal LRR or C-JID regions of the corresponding heteromeric receptor complexes (4–7). We identified multiple surface-exposed AVR_A6_, AVR_A10_, or AVR_A22_ residues required for recognition by the corresponding MLA receptors, indicating the existence of a similarly extensive effector– MLA receptor interface region. Together with the variations in the number, position, and length of individual structural elements found between AVR RALPH effectors, natural effector polymorphisms that locally affect individual structural AVR effector features essential for receptor recognition likely facilitate immunoevasion. Thus, our data support an evolutionary model in which changes in individual structural elements of RALPH effectors have contributed to the diversification of RALPH effector subfamilies, whereas nonstructural substitutions of surface-exposed effector residues have mainly driven diversification within a subfamily of RALPH effector paralogs.

The evolutionary model for RALPH effector diversification also provides a plausible framework to explain the molecular co-evolutionary arms race between MLA receptors and RALPH AVR_A_ effectors. The co-evolutionary dynamics of the barley–*Bgh* pathosystem must involve iterative cycles of generation and selection of novel MLA recognition specificities, followed by generation and selection of RALPH effectors that evade receptor detection to maintain pathogen virulence. Because *i*) AVR_A6_, AVR_A7_, AVR_A9_, AVR_A13_, and the allelic variants AVR_A10_/AVR_A22_ each belong to a different RALPH subfamily and are each detected by different MLA recognition specificities, *ii*) structural effector diversification has driven at least the differentiation of three subfamilies harboring AVR_A6_ or AVR_A7_ or AVR_A10_/AVR_A22_, and *iii*) these RALPH effector subfamilies also possess distinct conserved surface patches (**Fig. 5A**), it is likely that structure-driven RALPH subfamily differentiation is linked to the acquisition of distinct virulence functions. As a corollary, sequence-related RALPH effectors belonging to the same subfamily are probably targeting the same host process to promote virulence. Thus, we propose that proliferation of RALPH effectors in *B. graminis* genomes and their dominance in the effectorome was driven by both acquisition of novel virulence functions and escape from MLA receptor detection (**Fig. 5B**). This would also explain why multiple RALPH effector subfamilies and corresponding MLA recognition specificities are simultaneously maintained in host and pathogen populations.

How can allelic MLAs directly recognize effectors with divergent sequences? We identified six residues in AVR_A6_ that are essential for MLA6-dependent recognition (**Fig. 3; Fig. S6 & S7**). When we introduced these six residues (F27L, I31R, K33E, N36R, E40G and L47F) into CSEP0333, belonging to the subfamily containing AVR_A6_, the resulting construct was able to confer MLA6-dependent cell death in barley protoplasts and heterologous *N. benthamiana* (**Fig. 3**). However, when we constructed and tested a similar hybrid effector of AVR_A7_ containing the entire central segment of AVR_A6_, the resulting hybrid protein was highly unstable, and could often not be detected in *N. benthamiana* leaf extracts, indicating that structure-driven diversification of the two subfamilies containing AVR_A6_ or AVR_A7_ is connected to proper folding within subfamily members. Recently, crystal structures of two MAX effector variants APikL2F and APikL2A in complex with effector targets sHMA94 and sHMA25, respectively, revealed that a single polymorphic residue can lead to subtle changes in protein structures that ultimately determine the binding capacity of effectors to their host targets (47). Therefore, it is not surprising that structural elements of AVR_A_ effectors belonging to different subfamilies are not readily interchangeable. Allelic AVR_A10_ and AVR_A22_ were shown here to share a highly similar structure, but the residues involved in recognition by MLA10 and MLA22, respectively, locate to different surface patches on the subfamily-specific effector structure. Interestingly, MLA10 and MLA22 are not closely located within the phylogenetic tree of *Mla* resistance genes (48), which suggests that they evolved to detect AVR_A10_ and AVR_A22_, respectively, by convergent evolution. Therefore, MLA receptors have either evolved to detect distinct surface patches on RALPH effectors with highly similar structures or, as seems to be more common, surface patches of structurally diversified RALPHs belonging to different subfamilies. Furthermore, the structure of the wheat powdery mildew AVR_Pm2_ clarified here confirms that the phylogenetically unrelated CNL Pm2 has independently evolved the capability to recognize another site on RALPHs, termed the ‘head epitope’, in the sister species wheat, although whether this involves direct or indirect effector perception remains to be determined (24). It is likely that regions other than residues of RALPH effectors contacting the LRR contribute to MLA receptor activation. For instance, multiple contact residues in AvrSr35 are needed for binding to the Sr35 LRR domain, leading to a steric clash of other effector surface regions with the Sr35 NOD domain, with both of these processes needed for receptor activation (49).

The common fold adopted by *Blumeria* effectors raises the question of whether the conserved structural fold evolved convergently (i.e., from independent ancestral proteins) or whether the AVR RALPH effectors share a phylogenetic history which is masked by sequence diversification while the structure is retained (i.e., sequence-diversified from a common ancestor)? Previous work predicted that RALPHs share an intron in the same relative position (29, 41, 50), which points to a common ancestor. Indeed, in the five resolved RALPH AVR effector structures, an intron locates to the same relative position in the loop region between ß3 and ß4 (which is equivalent to ß2 and ß3 in AVR_A6_) (**Fig. S5**). Interestingly, this relative intron position is shared with RNase F1 and the catalytic active ribonuclease effectors from hemi-biotrophic Zt6 from *Zymoseptoria tritici*, Fg12 from *Fusarium graminearum* and SRE1 from *Setosphaeria turcica* (**Fig. S5**) (51–53). We conclude that RALPH AVR_As_ belonging to different subfamilies have evolved from an ancient ribonuclease that was present in the last common ancestor of Dothideomycetes and Sordariomycetes. Accordingly, maintenance of catalytic ribonuclease activity in the effectorome of *Z. tritici*, *F. graminearum* and *S. turcica* or its loss in powdery mildew, in tandem with neo-functionalization of multiple virulence functions, is associated with a transition in pathogen lifestyle from hemibiotrophic to obligate biotrophic.

## Materials and Methods

### Protein expression and purification

Effector sequences (22, 23, 25–29) were put into SignalP-5.0 Server Output - DTU Health Tech to detect signal peptides. The constructs used for effector protein purification have had the signal peptide removed. AVR_A1_(27–118), AVR_A6_(25–115), AVR_A7_(24–112), AVR_A9_(20–102), AVR_A10_(21–119), AVR_A13_(21–122), AVR_A22_(22–118), AVR_PM2_(21–119), AVRPM3^A2/F2^(24–130), AVRPM3^B2/C2^(21–130), AVRPM3^D3^(21–109), AVR_PM17_(24–110), AVRPM1a(18–155), AVRPM8(17–107), AvrSr50(24–132) and PWL2(21–145) were expressed in *E*. *coli* or insect cells as fusion proteins subcloned into pGEX-6P-1(GE Healthcare) or te pFastBac-1 vector (Invitrogen). These plasmids were used to express effectors with a N-terminal GST-tag followed by a PreScission proteolytic recognition site to remove the GST-tag.

Bacterial cultures were grown at 30 °C to an OD_600_ of around 0.8 in LB broth and induced with Isopropyl-β-D-thiogalactoside (IPTG, Sigma) for 15–18 h at 16 °C. The cells were harvested by centrifugation at 6,000 *g* for 10 min at 4 °C and resuspended in resuspension buffer (25 mM TRIS pH 8, 150 mM NaCl, 1 mM PMSF, 1 mM DTT). Bacterial cell suspensions were sonicated for 20 mins at 60% power (BANDELIN). Cell debris was removed by centrifugation at 30,000 *g* for 2 h at 4 °C. The soluble fractions were collected and allowed to flow through GST resin (GE Healthcare). After washing with two column volumes of the same buffer used for resuspension, another 2 ml of buffer and 10 μl of PreScission protease (GE Healthcare) were added to the column followed by overnight incubation to cleave off the AVR proteins from the GST resin. The cleaved AVR proteins were then eluted and further purified by size-exclusion chromatography using a HiLoad 16/600 Superdex 200 pg gel filtration column (GE Healthcare). Purified proteins were concentrated to 10–30 mg/ml by using a 10-kDa Amicon centrifugal filter device (Merck), flash-frozen in liquid nitrogen, and stored at −80 °C. Baculoviruses (50 ml) for AVR expression were individually added to 1 L of SF21 insect cells (1.8 x 10^6^ cell ml^-1^) cultured at 28 °C in Sf-900 II SFM medium. The medium was collected 48 h after infection. The purification process is the same as for the *E. coli* system.

### Crystallization, data collection, structure determination and refinement

The initial crystallization experiments were carried out at 20 °C, using the sitting-drop vapor-diffusion method. For screening, the AVR effector proteins were mixed 1:2, 1:1 and 2:1 with different crystallization buffers using a Mosquito Nanodrop. Out of 16 effector proteins, good diffraction crystals were obtained for only AVR_A6_, AVR_A7_, AVR_A10_, AVR_A22_, AVR_PM2_. After the initial screening, further optimization was performed using a 24-well hanging-drop vapor-diffusion method with an equal volume (1.0 µl) of protein and reservoir solution at 20 °C. Crystals with the best morphology were observed in 20 % w/v polyethylene glycol 3 350, 200 mM sodium fluoride for AVR_A6_; 1.4 M sodium phosphate monobasic monohydrate/potassium phosphate dibasic pH 8.2 for AVR_A7_; 0.16 M calcium acetate hydrate, 0.08 M sodium cacodylate trihydrate pH 6.5, 14.4% w/v polyethylene glycol 8,000 for AVR_A10_; 1.0 M succinic acid pH 7.0, 0.1 M HEPES pH 7.0, 1% w/v polyethylene glycol monomethyl ether 2,000 for AVR_A22_; 0.1 M BIS-TRIS pH 6.5, 28% w/v polyethylene glycol monomethyl ether 2,000 for AVR_PM2_. Crystals were transferred into a cryoprotectant solution containing a reservoir solution with 20% glycerol. The diffraction data were collected at different beamlines as indicated in the **Table S1**. The data were processed using XDS or autoProc (54, 55). The crystal structures of these five AVR effectors were determined by molecular replacement (MR) with Phenix using structures predicted by AF2 as the initial search model. The models from MR were built automatically by ModelCraft (56) and/or computer-assisted with COOT (57) and subsequently subjected to refinement by Phelix software suite (58). Statistics of diffraction data and refinement of these five effector models are summarized in **Table S1**. Structural figures were prepared using the program ChimeraX v1.3 (59). Sequence alignments were processed with the ENDscript server (60).

### AVR_A_ effector RNase activity assays

Total RNA was extracted from 9-day-old barley cv. Golden promise plants using the RNeasy Plant Mini Kit (QIAGEN) and treated with TURBO DNase enzyme (Ambion) to remove genomic DNA. Purified AVR_A_ effectors from *E. coli* were then incubated with the total barley RNA. The reaction mixture consisted of 1 μg of RNA and 1 μM of protein and was prepared in a buffer containing 15 mM Tris-HCl (pH 8.0), 15 mM NaCl, 50 mM KCl, and 2.5 mM EDTA. The reaction mixture was incubated at 25 °C for 90 minutes. For analysis using the Bioanalyzer 2100 (Agilent Technologies, USA), 10 μl of the sample were used. RNase T1 (Thermo Scientific) was included as a positive control in the assay.

### Production and detection of 2’, 3’-cNMP *in vitro*

Barley total RNA (100 ng) was individually incubated with purified AVR effector (1 µM for each), L7^TIR^ (1 µM), and 2.5 µl of RNase T1 (Thermo Scientific) in buffer containing 25 mM Tris-HCl pH 8.0 and 150 mM NaCl at 25 °C for 16 h. The total volume for each reaction was 50 µl. The samples were centrifuged at 12, 000 *g* for 10 min and the supernatant was applied to LC-MS/MS for metabolite measurement.

### Metabolite measurement by LC-MS/MS

Chromatography was performed on a Nexera XR 40 series HPLC (Shimadzu) using a Synergi 4 μM Fusion-RP 80 Å 150×2 mm column (Phenomenex). The method of determination of 2’3’cNMP is described in (43).

### Transient gene expression assays in barley protoplasts

Entry clones and destination constructs for the expression of *AVR_A6_, AVR_A7_, MLA6 and MLA7* were previously published by (22, 29). Entry clones for *CSEP0333* (BLGH_00698), the chimeric effectors A6N, A6M, A6C, B6N, B6M, B6C, M1 and M2 were generated by gene synthesis based on wild-type codons (GeneArt, Invitrogen). The constructs M1 and M2 were used as templates to generate higher-order mutants by site-directed mutagenesis PCR (NEB, Q5 Site-Directed Mutagenesis Kit) using the primers listed in Table S2. The integrity of all entry clones was confirmed by Sanger sequencing. For transient expression assays in barley protoplasts and *N*. *benthamiana* leaves, the genes were recombined using LR-Clonase II (Thermo Fisher) into the pIPKb002 (*Spec^R^*) (61) gateway-compatible destination vectors. The integrity of all expression vectors was confirmed by Sanger sequencing. The isolation and transfection of barley leaf protoplasts was performed as described in (44). cDNAs of the *AVR_a_* effectors chimeras were co-expressed with cDNAs of *MLA6 or MLA7* using the pIPKb002 vector with the ubiquitin promoter in barley *cv*. Golden Promise protoplasts. Protoplast solution (300 μl of 3.5 x 10^5^ cells ml^−1^) was transfected with 4 μg of *LUC* reporter construct (pZmUBQ: LUC), 12 μg of *Mla* plasmid, and 5 μg of the respective *AVR_a_* (chimeric) effector or an empty vector (*EV*).

### Transient gene expression in *N. benthamiana* and protein detection by immunoblotting

For *N. benthamiana* transient gene expression, *AVR_A6_, CSEP0333* and effector chimeras and mutants were cloned into the *pDONR221* vector (Invitrogen). The obtained plasmids were recombined by an LR clonase II (Thermo Fisher Scientific) into the *pXCSG-mYFP* vector with a C-terminally fused mYFP epitope tag. Constructs were verified by Sanger sequencing. The *Mla6* and *Mla7* expression clones were previously described by (22, 29). Expression constructs were transformed into *Agrobacterium tumefaciens* GV3101 (pMP90RK) by electroporation. Transformants were grown on LB media selection plates containing rifampicin (15 mg ml^−1^), gentamycin (25 mg ml^−1^), kanamycin (50 mg ml^− 1^), and spectinomycin (50 mg ml^−1^) for transformants harboring *pGWB517-Mla6-4×Myc* or carbenicillin (50 mg ml^−1^) (62) for *pXCSG-mYFP* effector constructs (63).

Individual *Agrobacterium* transformants were cultured in LB medium containing respective antibiotics at 28 °C for 16 h. Bacterial cells were harvested by 2500 *g* for 15 min and resuspended with infiltration buffer containing 10 mM MES pH 5.6, 10 mM MgCl_2_ and 150 μM acetosyringone. Construct expression was conducted in leaves of four-week-old *N. benthamiana* plants via *Agrobacterium*-mediated transient expression assays in the presence of the P19 and CMV2b suppressors of RNAi silencing (64). The final OD_600_ of receptor, effector and RNAi silencing suppressor strains was adjusted to 0.5 each. Phenotypic data were recorded at day 6 after infiltration. For protein detection, the leaf material from four individual plants was harvested 48 h after infiltration, flash-frozen in liquid nitrogen and ground to powder using a Retsch bead beater. Plant powder was mixed with 4 x Laemmli buffer in a 1:2 ratio. After centrifugation at 16,000 *g* for 15 min, 5 µl of supernatant were loaded onto a 10% SDS-PAGE. Separated proteins were transferred to a PVDF membrane and probed with monoclonal mouse anti-Myc (1:3,000; R950-25, Thermofisher), polyclonal rabbit anti-GFP (1:3,000; pabg1, Chromotek) followed by polyclonal goat anti-mouse IgG-HRP (1:7,500; ab6728, Abcam) or polyclonal swine anti-rabbit IgG-HRP (1:5,000; PO399, Agilent DAKO) antibodies. Protein was detected using SuperSignal West Femto: SuperSignal substrates (ThermoFisher Scientific) in a 1:1 ratio.

### Microscale Thermophoresis (MST)

For Microscale Thermophoresis experiments, total RNA was isolated from 7-day-old barley cv. Golden promise plants by phenol/chloroform extraction. Briefly, 5 g of leaf material was ground to a fine powder in liquid nitrogen. In a 50 mL propylene tube, the powder was resuspended in 10 mL lysis buffer (100 mM TRIS pH 8.0, 100 mM NaCl, 20 mM EGTA, 2% SDS) and 100 µL 2-Mercaptoethanol, followed by the addition of 1 volume of Phenol. Tubes were incubated for 20 min while mixing in a revolving rotator, followed by the addition of 0.5 volume mL Chloroform and another 15 min mixing. Samples were centrifuged for 10 min at maximum speed, and the upper aqueous phase was transferred to a fresh tube. Phenol/Chloroform extraction was repeated total of three times, followed a fourth time with Chloroform only. Then, nucleic acids were precipitated by the addition of 0.1 volumes of DEPC-treated 3M Sodium Acetate pH 5.2 and 2.5 volumes of Ethanol, following by incubation at −70 °C for >30 min. After centrifugation at 30 min at max. speed, pellets were resuspended in 5 mL of DEPC-treated water, followed by addition of 5 mL DEPC-treated LiCl and incubation on ice at 4 °C for >3 hrs. Finally, after centrifugation at 30 min at max. speed, pellets were resuspended in 1.8 mL DEPC-treated water and precipitated one more time using Sodium Acetate and Ethanol, followed by three washing steps with 70% Ethanol. RNA pellets were resuspended in 500 µL DEPC-treated water. To obtain RNA concentrations >5 µg µL^-1^, the RNA pellets from 24 extractions were pooled.

The fluorescent dye NT-647 (MO-L001, NanoTemper Technologies) was used to label effector proteins GST or BSA. The labeled proteins were eluted with the reaction buffer (20 mM phosphate-buffered saline, 150 mM NaCl, and 0.05% (v/v) Tween 20, pH 7.4), and mixed with different concentrations of barley total RNA (Phenol/Chloroform) before loading onto Monolith NT.115 (NanoTemper Technologies). Data were treated by the KD Fit function of the Nano Temper Analysis Software (version 1.5.3).

### Phylogenetic analysis of RALPH effectors and detection of conserved surface-exposed amino acids

A maximum likelihood phylogeny was constructed according to (29), including all predicted CSEPs from *B. graminis* f sp *poae*, *lolium*, *avenae*, *tritici* 96224, *hordei* DH14, *secalis* S1459, *triticale* T1-20, and *dactylidis*. The protein sequences of the members of effector subfamilies were aligned using MUSCLE and then displayed by ESPript3 (https://espript.ibcp.fr/ESPript/ESPript/). Conserved residues >70% have been highlighted in the crystal structures.

## Acknowledgements

We thank the Alexander von Humboldt Foundation (J.C.), the Max-Planck-Gesellschaft (P.S.-L. and J.C.), the Deutsche Forschungsgemeinschaft (DFG, German Research Foundation) in the Collaborative Research Centre Grant (SFB-1403 – 414786233 B08 to J.C. and P.S.-L.) and Germany’s Excellence Strategy CEPLAS (EXC-2048/1, Project 390686111; J.C. and P.S.-L.) for funding of this project. Y.C. was funded by a PhD fellowship from the Chinese Scholarship Council (number 201808440401). Crystals were grown in the Cologne Crystallisation facility (http://C2f.uni-koeln.de) supported by the DFG (Grant No. INST 216/682-1 FUGG). We thank the staff of the beamlines X06SA (PXI, PSI, Switzerland), EMBL Hamburg at the PETRA III storage ring (DESY, Germany) and ESRF (Grenoble, France) for their help during data collection. AVR_A22_ and AVR_PM2_ data were collected at X06SA, AVR_A6_ and AVR_A7_ at beamlines P13 (EMBL; (65) ) and P14 (Proposal MX828) and AVR_A7_ at ID30B (ESRF; (66) Proposal MX2412). We thank Gleb Bourenkov and Saravanan Panneerselvam for assistance in using these beamlines, Neysan Donnelly for manuscript editing and Petra Köchner for assistance with recombinant DNA work.

## Author Contributions

J.C., and P.S.-L. conceived the study; Y.C., F.K., J.C., and P.S.-L. designed experiments; Y.C., F.K., E.L., J.M.G., A.W.L., D.Y. and J.J. performed research; Y.C., F.K., J.M.G., U.B., M.U. and K.T. analyzed data; B.K. contributed new reagents/analytic tools. Y.C., F.K., J.C., and P.S.-L. wrote the paper with input from all authors.

## Competing Interest Statement

The authors declare no competing interests.

## Material, data, and code availability

All study data are included in the article and/or supporting information. Data deposition: The atomic coordinates have been deposited in the Protein Data Bank, www.pdb.org [PDB ID codes 8OXH (AVR_A6_), 8OXL(AVR_A7_), 8OXK (AVR_A10_), 8OXJ (AVR_A22_) and 8OXI(AVR_PM2_)].

## Supplementary figures

**Figure S1.**
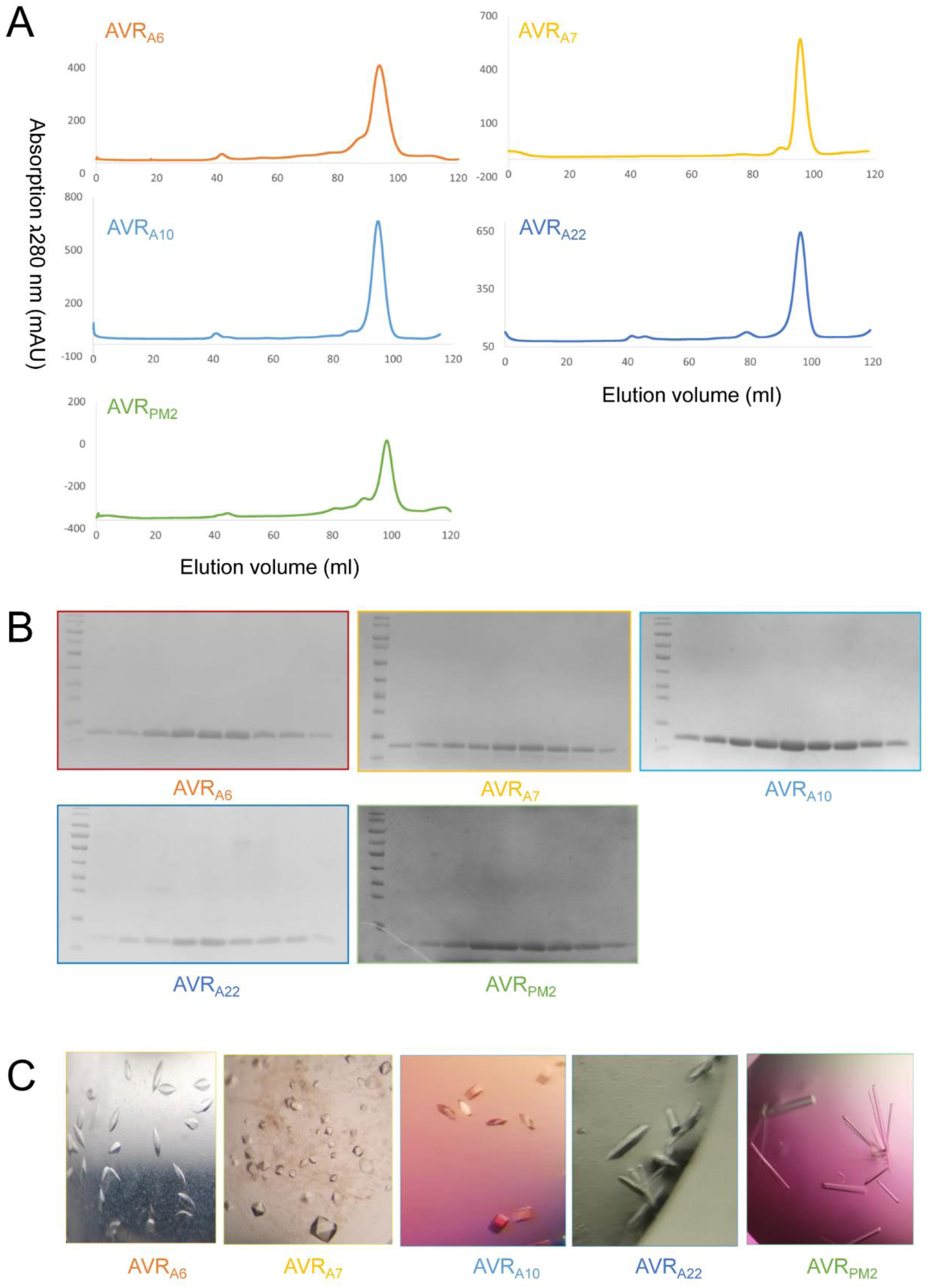
Purification and crystallization of *Blumeria graminis* AVR effector proteins. (A) Absorption spectra at 280 nm (mAU) for the effectors that were purified using Size-exclusion chromatography (SEC). (B) Coomassie staining of selected peak fractions. (C) Representative pictures of crystals obtained for five AVR effectors.

**Figure S2.**
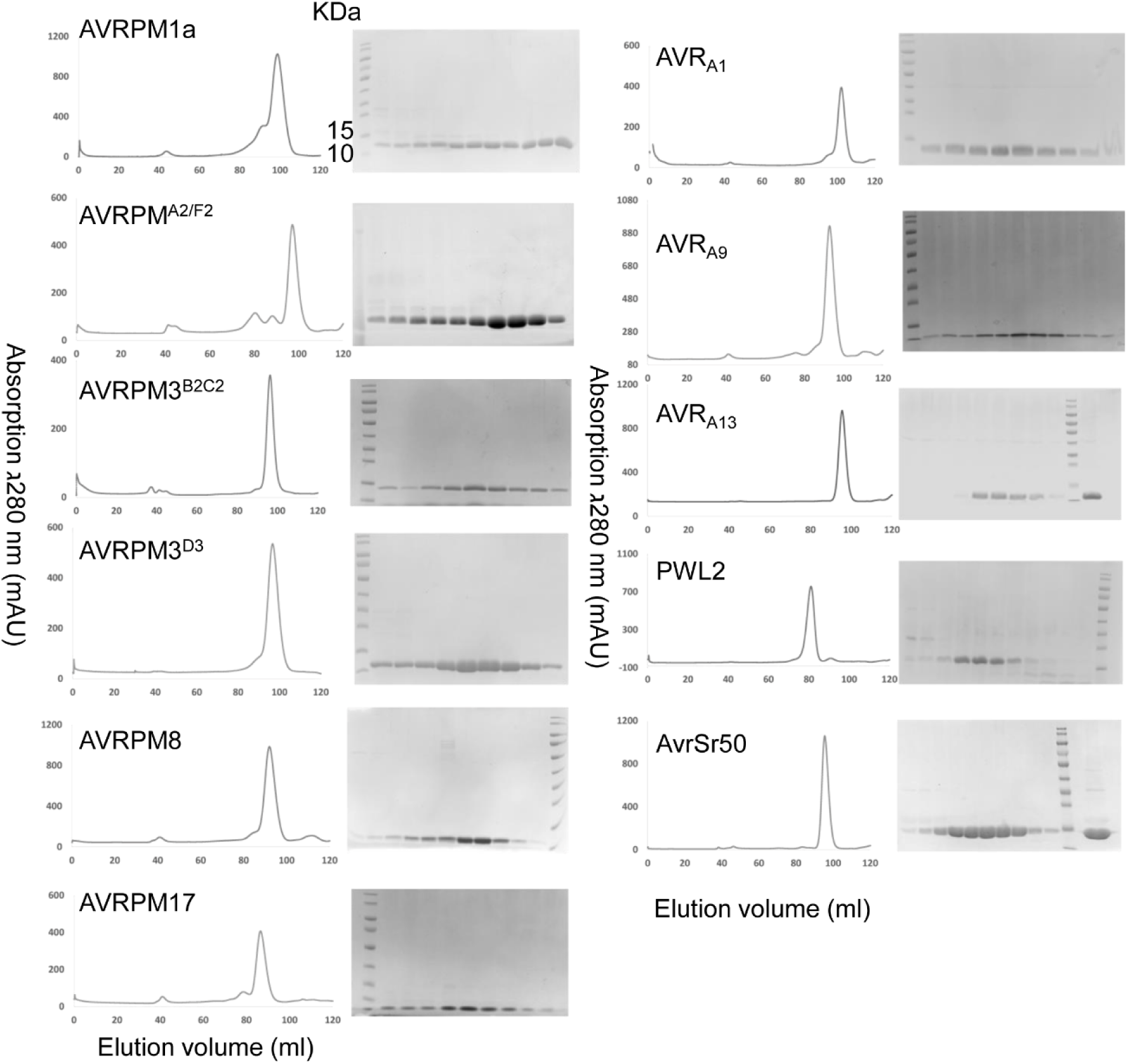
Purification of additional ascomycete effectors. 11 additional effectors were purified to homogeneity using SEC but failed to subsequently yield well-diffracting crystals.

**Figure S3.**
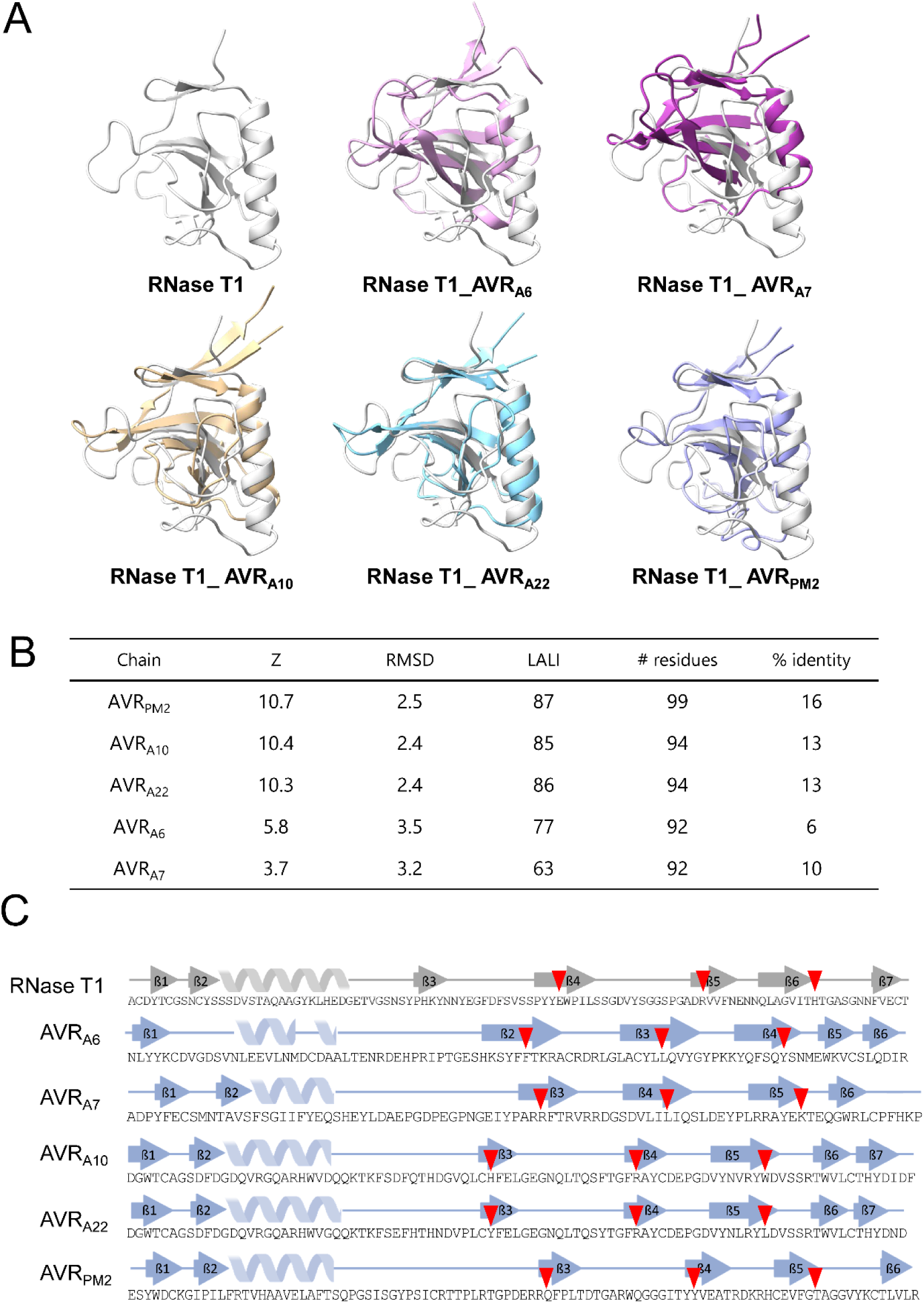
Structural comparison of *Blumeria graminis* AVR effectors with RNase T1 from *Aspergillus oryzae*. (A) Superimposition of the AVR structures with RNase T1 (PDB: 9RNT) in cartoon representation. (B) Pairwise structural comparisons using the DALI server. (C) 2D-representation of the RNase T1 structure and AVR effectors with residues corresponding to the catalytic triad in RNase T1 highlighted with a red triangle.

**Figure S4.**
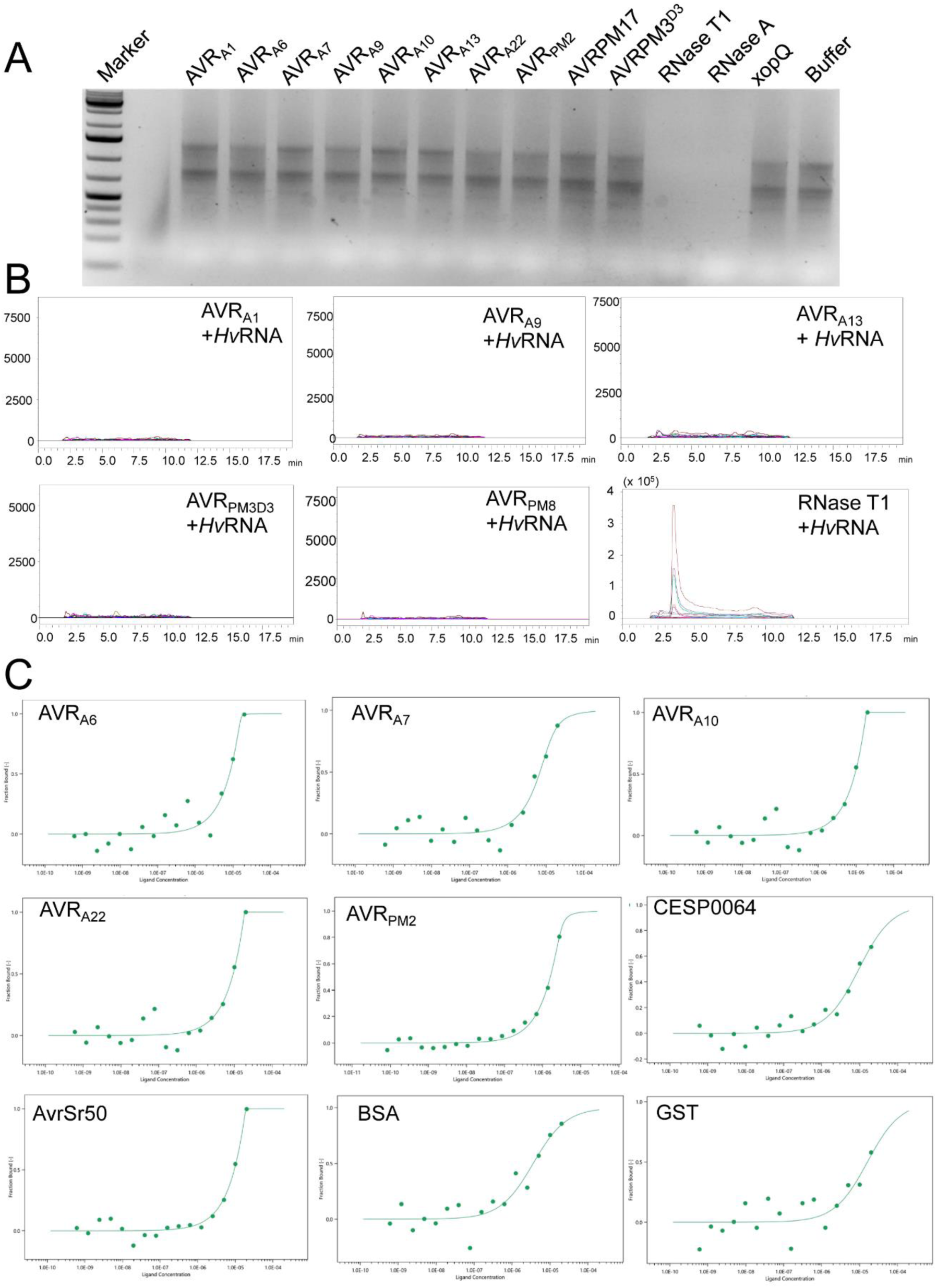
*Blumeria graminis* AVR effectors are pseudo RNases. (A) Co-incubation of RNA with AVR effectors for 16 hours and subsequent gel electrophoresis (1% agarose gel, 100V, 45 min) Marker: GeneRuler 1kb Plus DNA ladder (Invitrogen). (B) Detection of 2’,3’-cNMP synthetase activity using LC-MS for additional AVR effector proteins. (C) MST traces of AVR effectors as well as non-RNase-like fold proteins BSA, GST and AvrSr50 with *Hv*RNA. All proteins were recombinantly expressed, purified and subsequently labelled using the MO-L001 labelling kit (NanoTemper). The highest RNA concentration was set to 3750 ng/µL.

**Figure S5.**
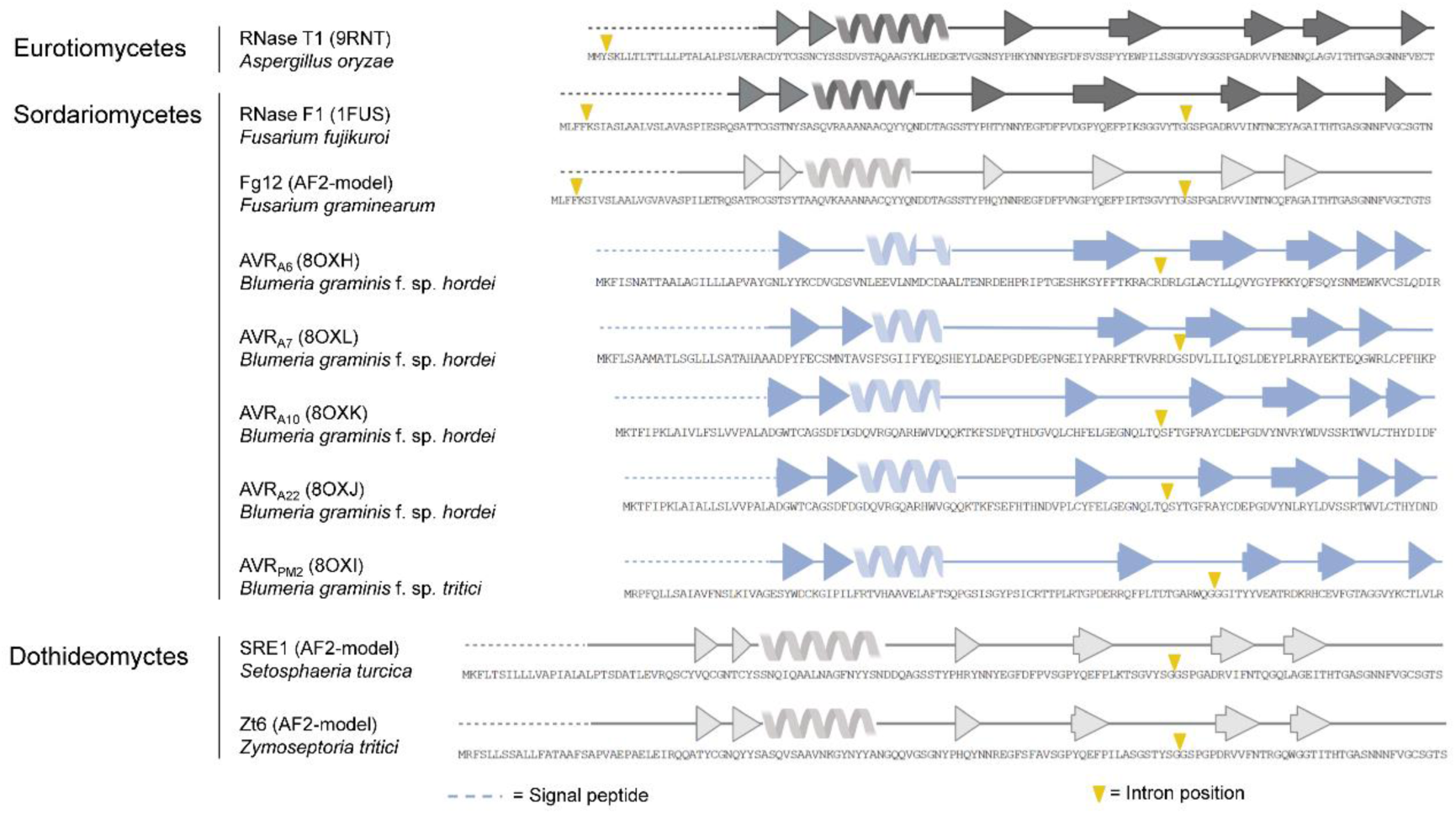
*Blumeria graminis* AVR effectors share a single intron with RNase F1 family members. 2D-representation of structures or AlphaFold2-models of characterized secreted ribonucleases and RALPH effectors with intron positions in the translated sequences highlighted with a yellow triangle. Signal peptide sequences are pictured with a dashed line. NCBI gene identifiers used: RNase T1: AP007171.1; RNase F1: AB355898.1; Fg12: FG11190.1; Zt6: NC_018216.1; SRE1: NW_007360025.1 (SETTUDRAFT_163271); AVR_A6_: UNSH01000074 (BLGHR1_15960); AVR_A7_: UNSH01000097 (BLGHR1_17217) AVR_A10_ and AVR_A22_: CAUH01000387.1 (BGHDH14_bgh03730) AVR_PM2_: KX765276.1.

**Figure S6.**
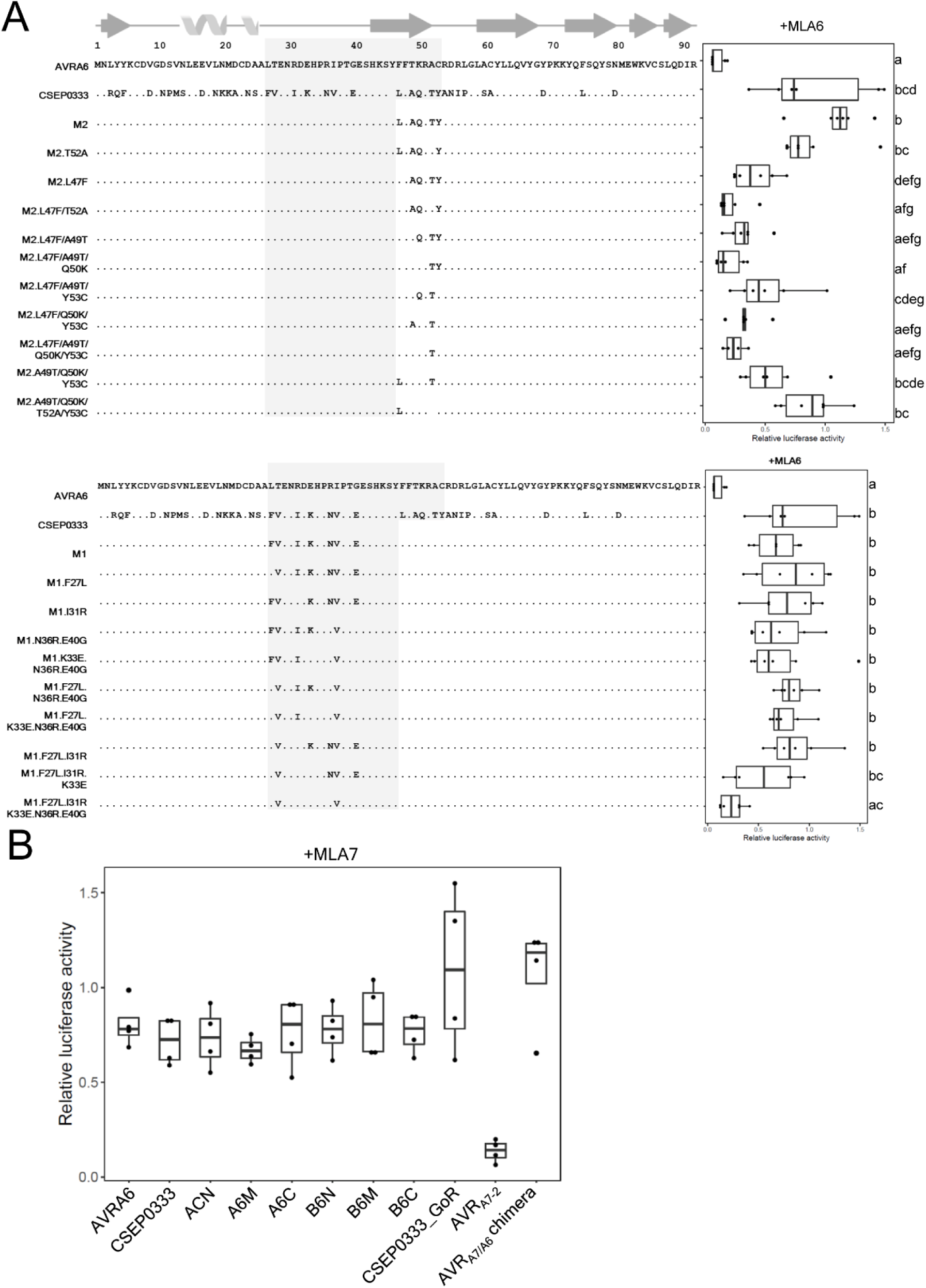
Six amino acids in the central segment of AVR_A6_ are essential for the detection by MLA6 in barley protoplast. (A) Hybrid effectors with targeted substitutions were co-transfected with MLA6 into barley protoplasts and cell death was quantified by measuring luciferase reporter activity. Letters indicate results of statistical variance analysis using Kruskal-Wallis test followed by Dunn’s post hoc tests (P < 0.05). Raw relative luciferase measurements and P-values for all protoplast plots are provided in Supplementary Data S6. (B) Selected effector constructs were co-transfected with MLA7 into barley protoplasts as a control for receptor specificity.

**Figure S7.**
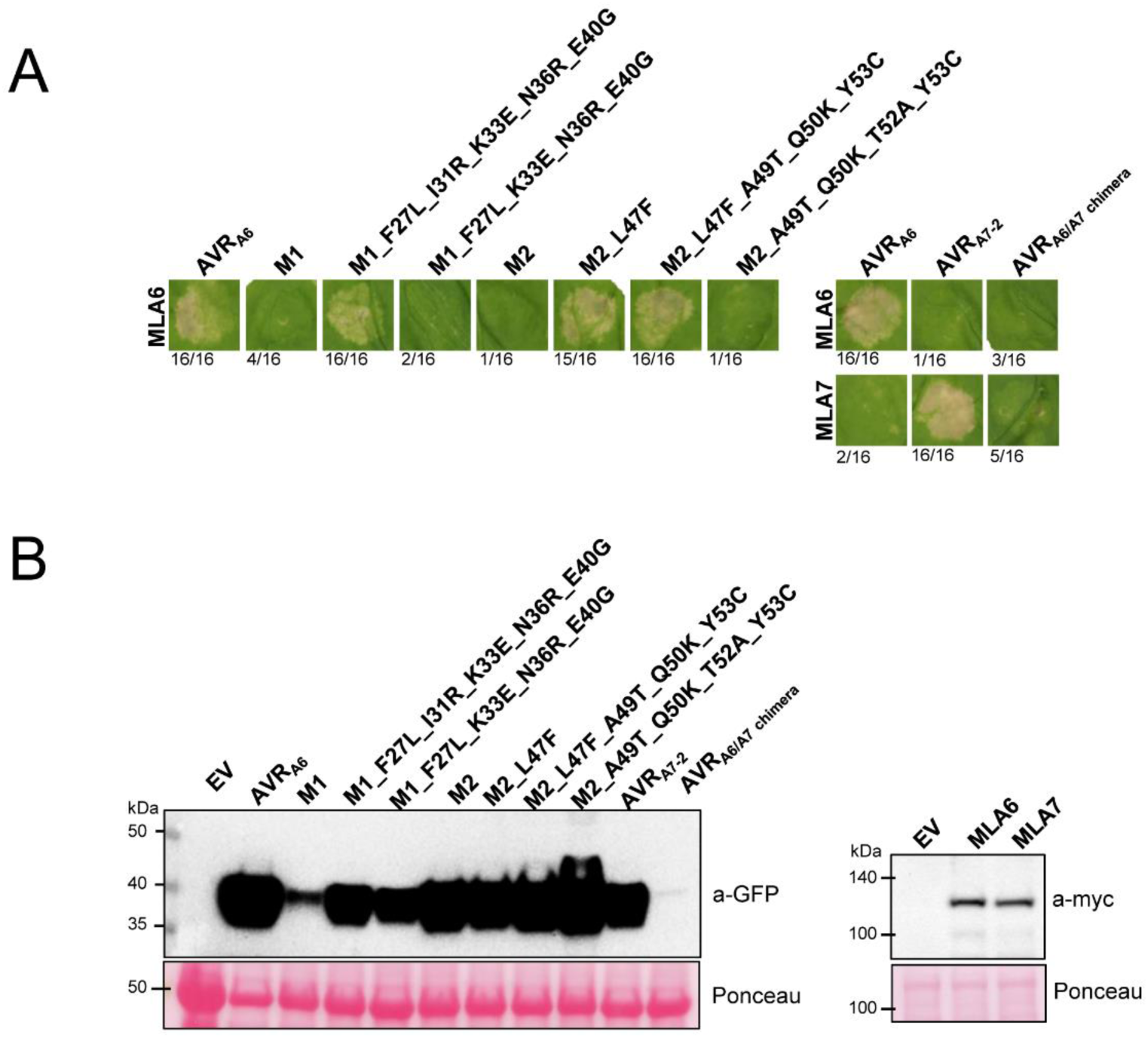
Six amino acids in the central segment of AVR_A6_ are essential for the detection by MLA6 in *N. benthamiana*. (A) Selected effector hybrids with targeted amino acid substitutions were co-expressed in *N. benthamiana* using *Agrobacterium*-mediated infiltration. The cell death score is indicated below the representative pictures from 16 replicates. (B) Immunoblot to detect accumulation of effector and receptor proteins.

**Figure S8.**
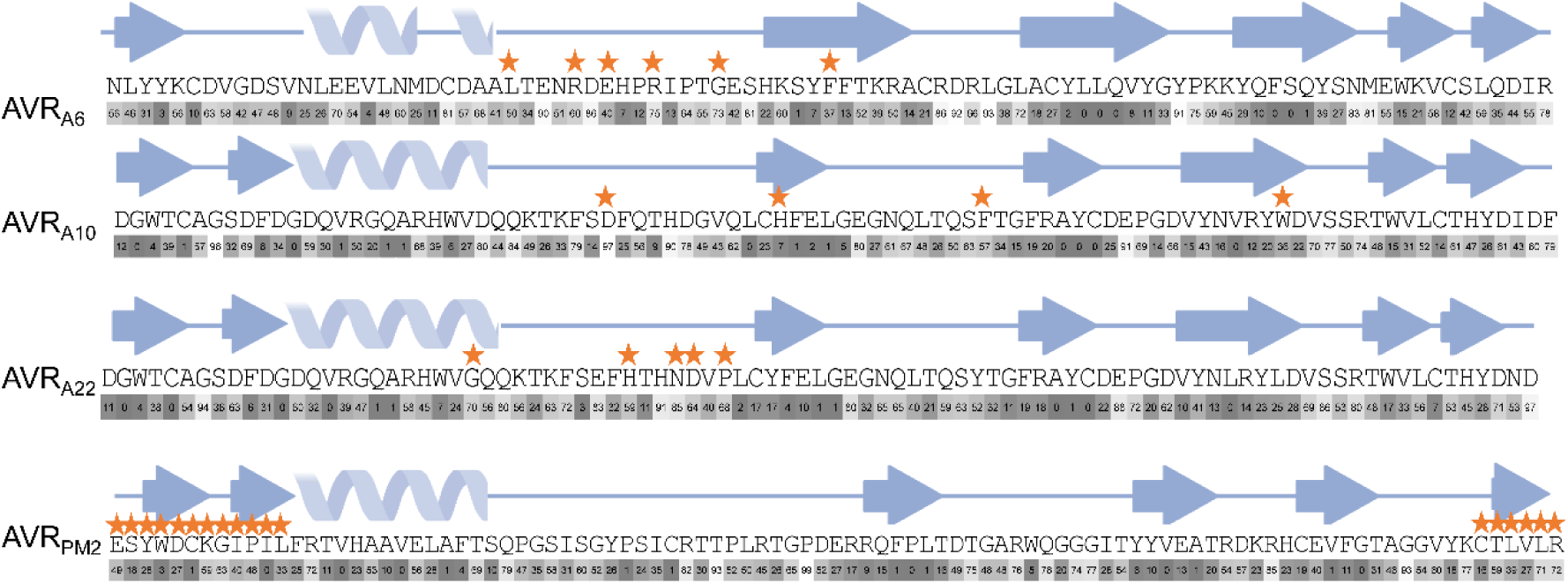
Relative solvent-accessibility of *Blumeria graminis* AVR effector sequences. Relative solvent-accessibility was computed using PyMOL. Orange asterisk indicate the residues important for recognition by the cognate NLR receptor.

**Figure S9.**
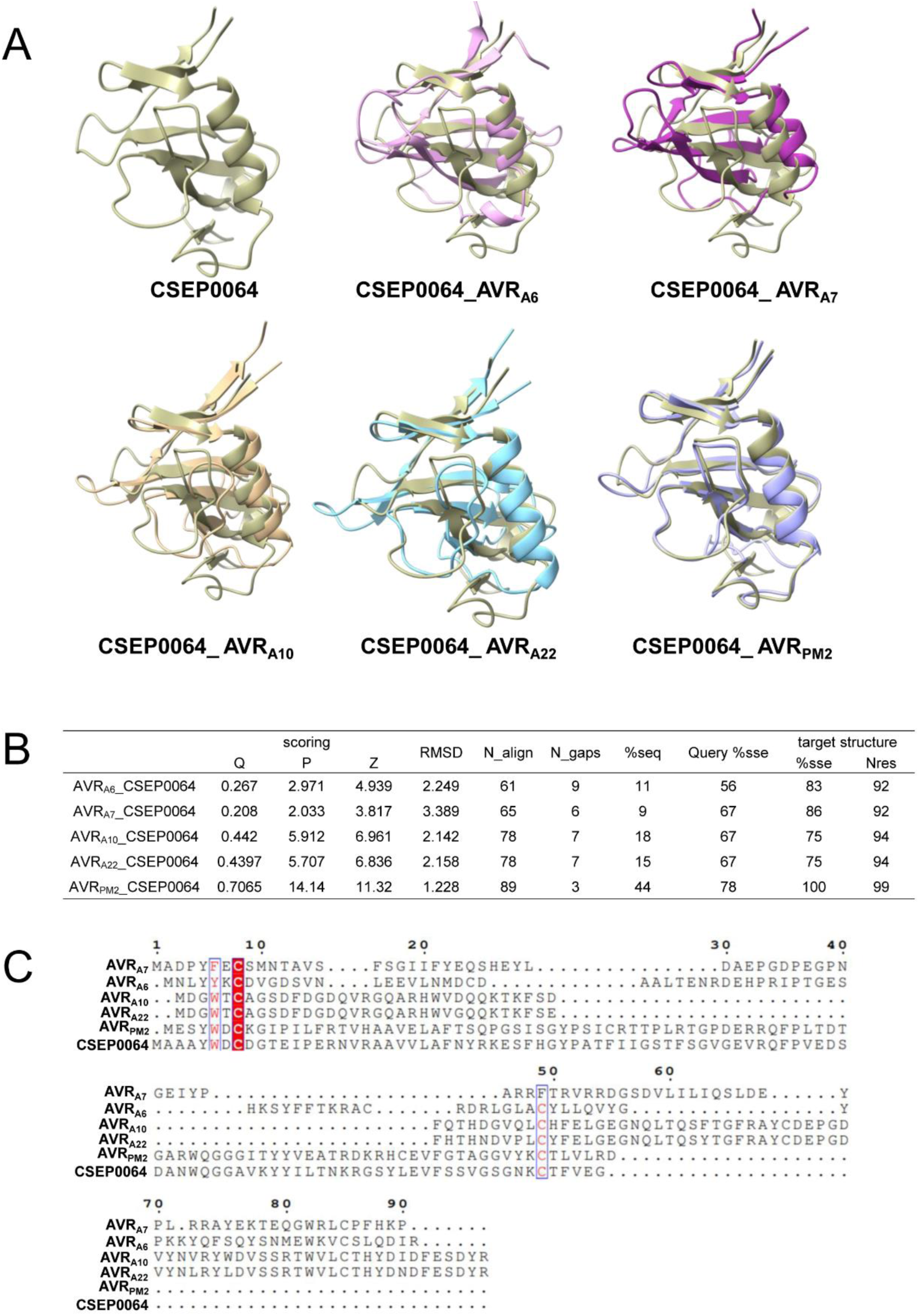
Structural comparison of *Blumeria graminis* AVR effectors with CSEP0064. (A) Superimposition of the AVR structures with CSEP0064 (BEC1054) (PDB: 6FMB) in cartoon representation. (B) Pairwise structural comparisons using the DALI server. (C) Sequence alignment of AVR structures with CSEP0064.

**Figure S10.**
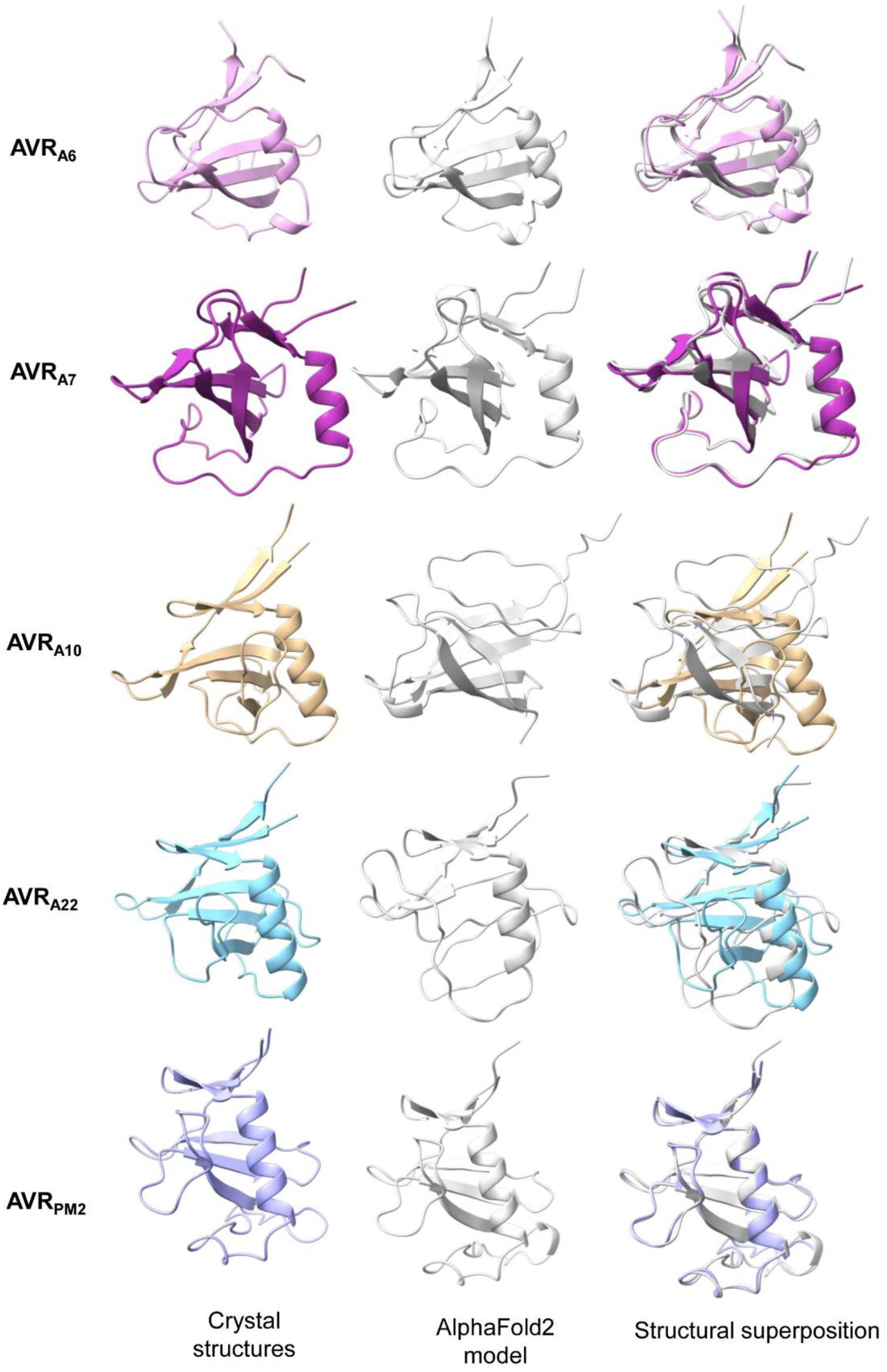
Structural comparison of the AlphaFold2 predicted and experimentally determined structure of AVR effectors. (A) For structural modelling, Colabfold v1.3 (colab.research.google.com/github/sokrypton/ColabFold/blob/main/AlphaFold2.ipynb) was used to predict the structures of the AVR effector without their signal peptide. The top rank model was used for superimposition with the experimental structure.

**Table S1.**
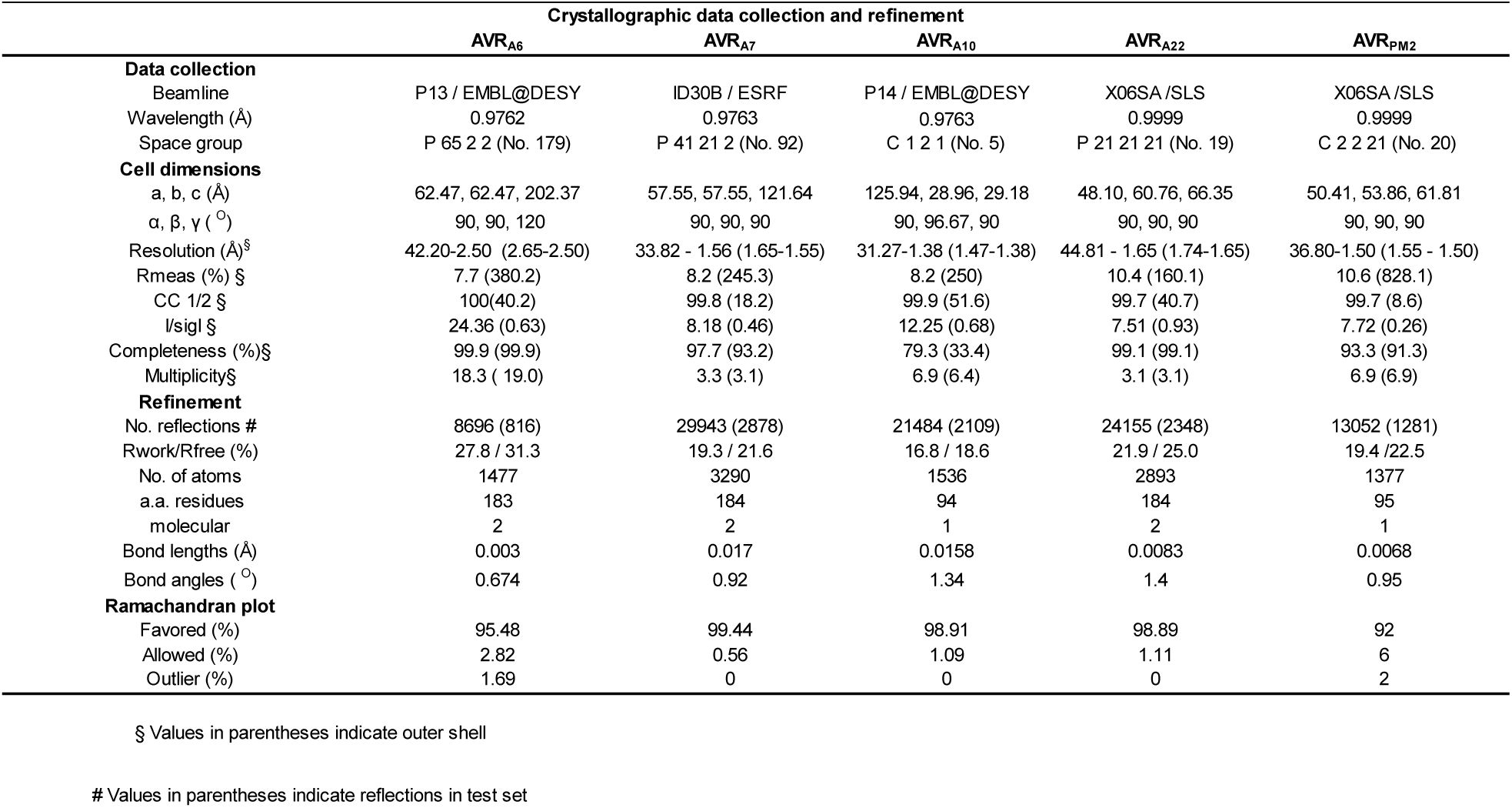
Crystallographic data collection and refinement.

**Table S2.**
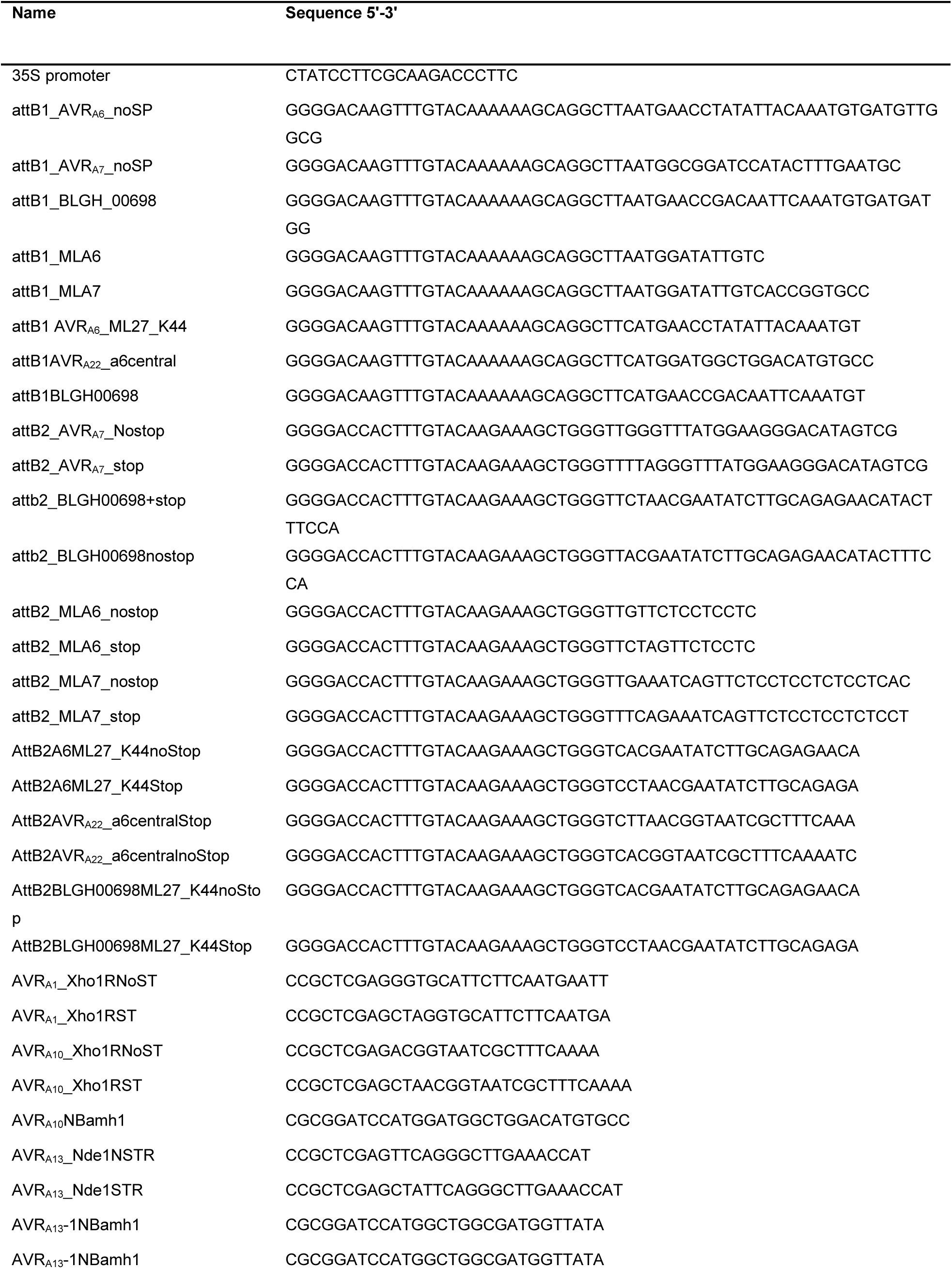
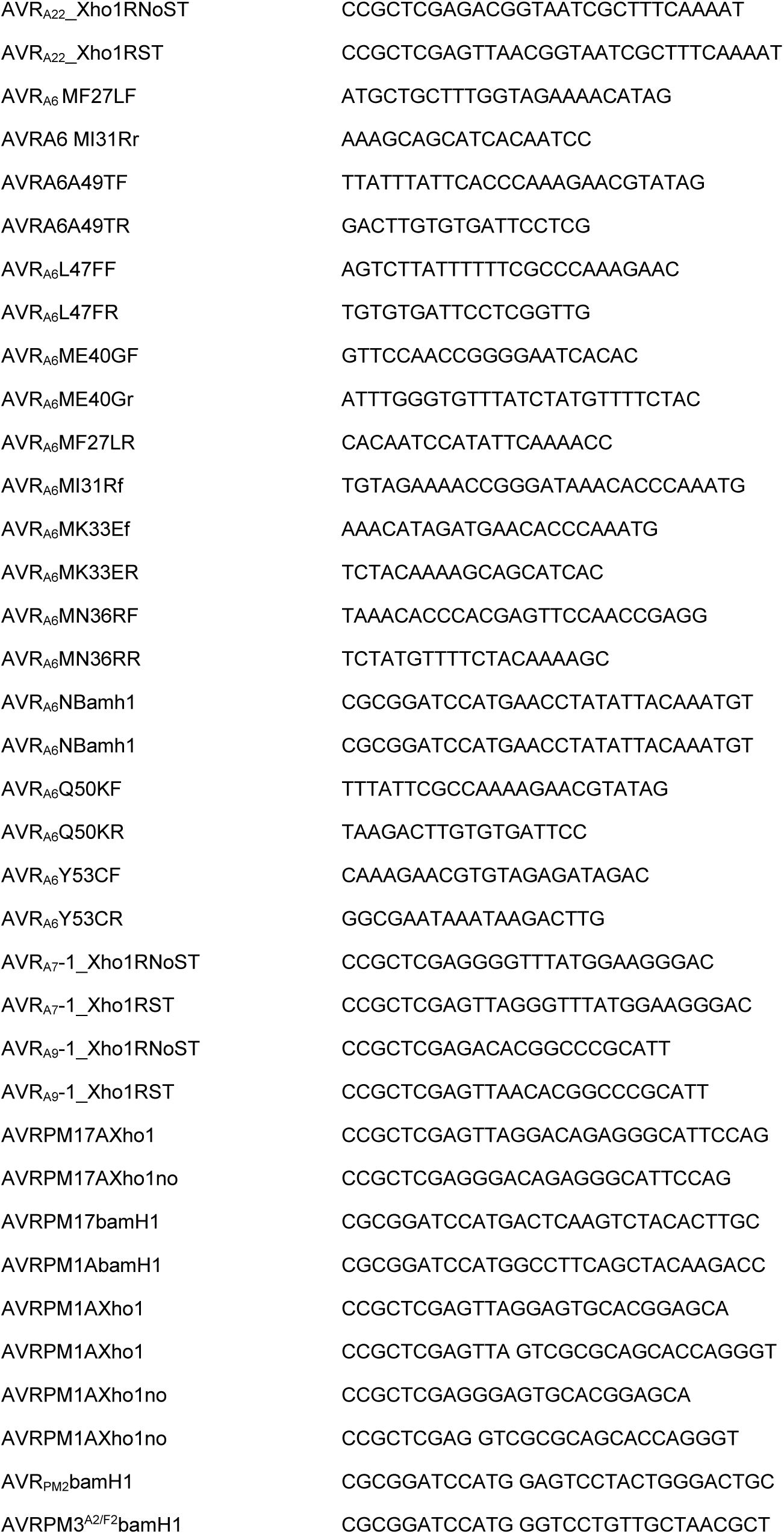
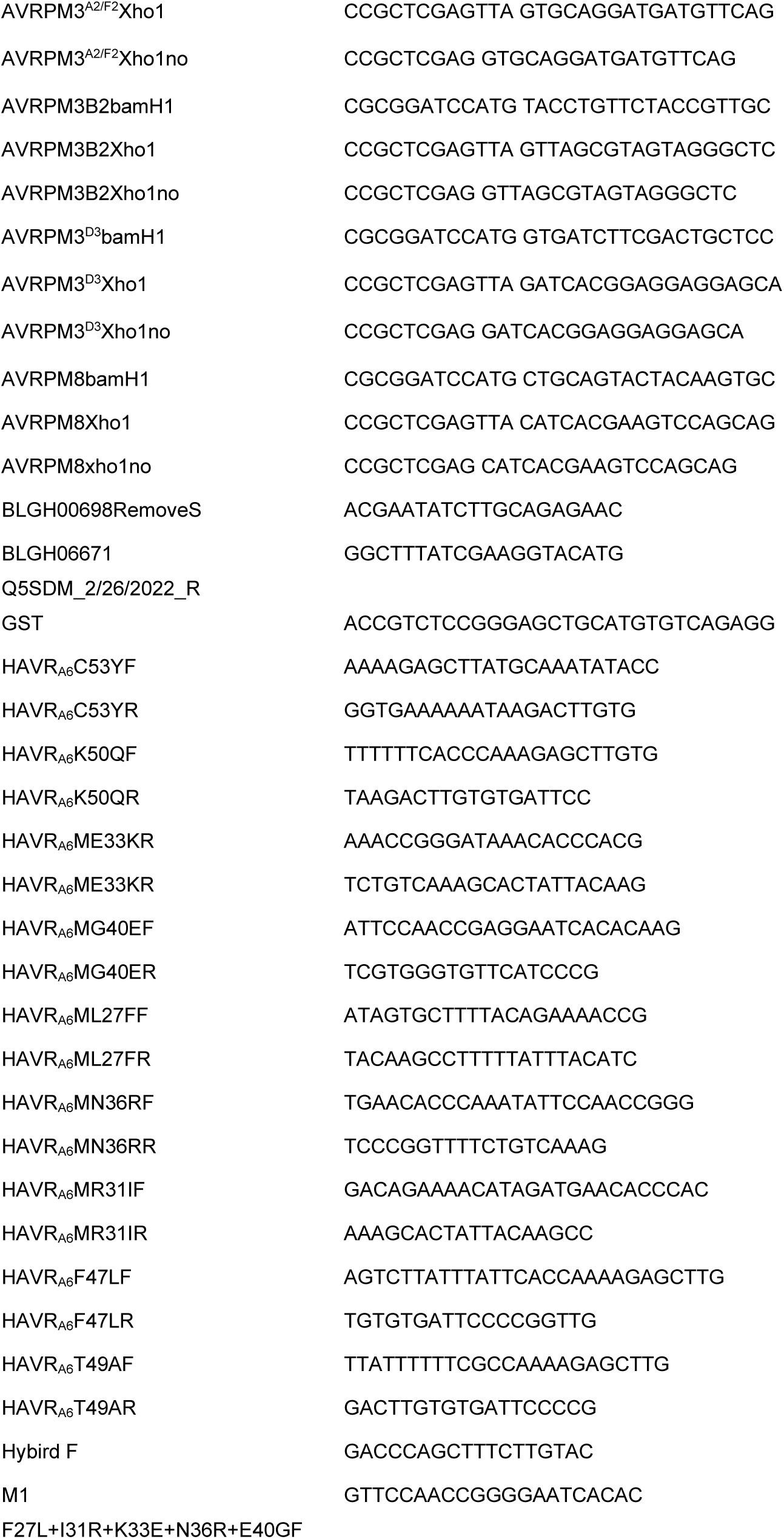
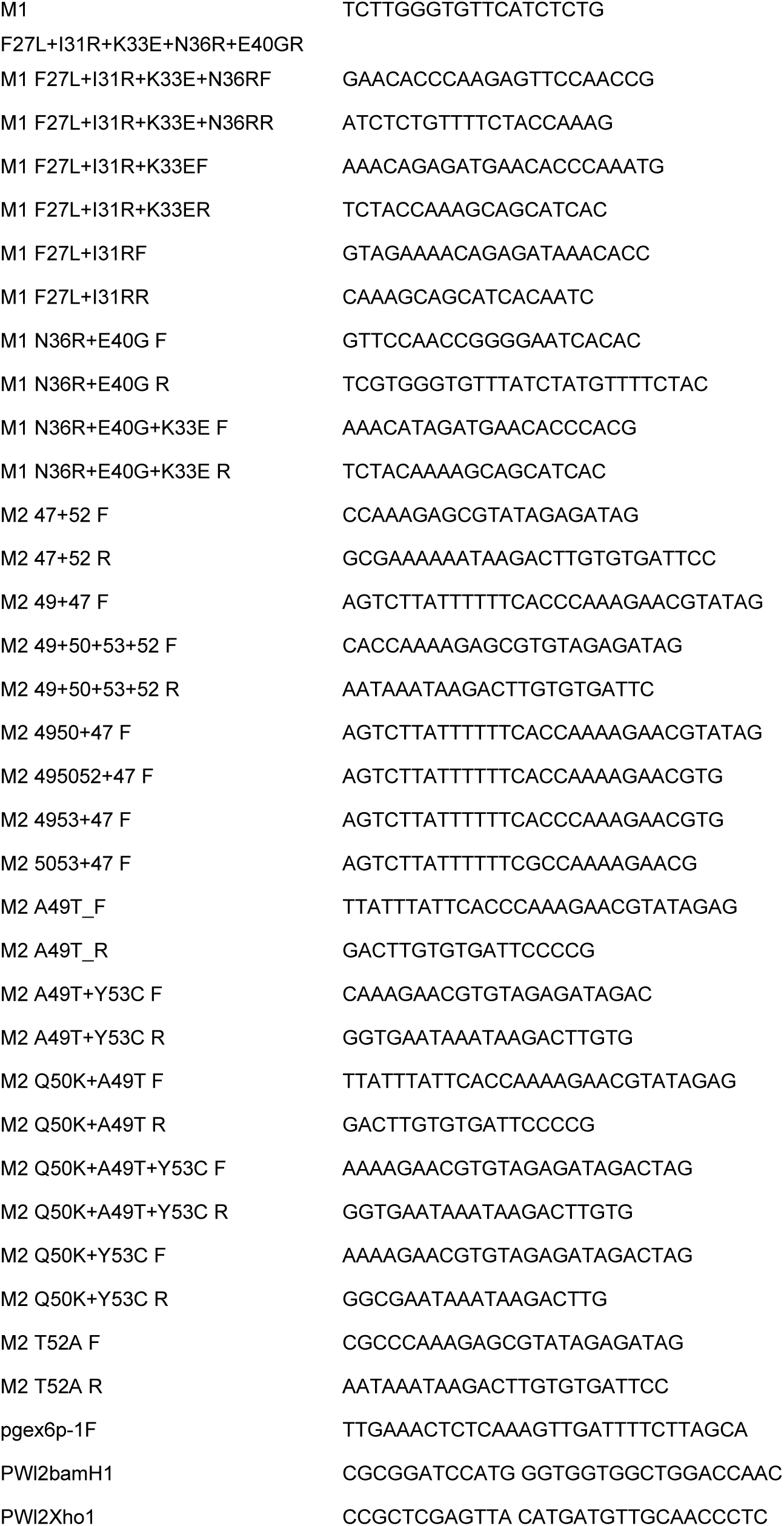
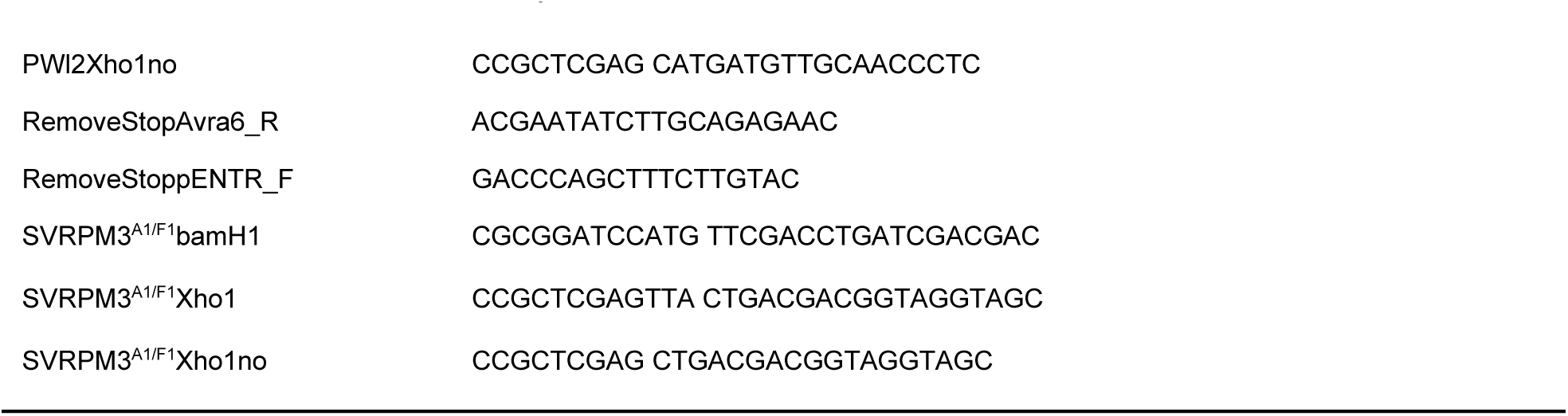
Primers used in this study.

## Notes

### Competing Interest Statement

The authors have declared no competing interest.

## References

1. I. M. Saur, R. Panstruga, P. Schulze-Lefert, NOD-like receptor-mediated plant immunity: from structure to cell death. Nat Rev Immunol 21, 305–318 (2021).

2. Z. Hu, J. Chai, Assembly and Architecture of NLR Resistosomes and Inflammasomes. Annual Review of Biophysics 52, (2023).

3. D. Lapin, O. Johanndrees, Z. Wu, X. Li, J. E. Parker, Molecular innovations in plant TIR-based immunity signaling. The Plant Cell 34, 1479–1496 (2022).

4. S. Ma et al., Direct pathogen-induced assembly of an NLR immune receptor complex to form a holoenzyme. Science 370, eabe3069 (2020).

5. R. Martin et al., Structure of the activated ROQ1 resistosome directly recognizing the pathogen effector XopQ. Science 370, eabd9993 (2020).

6. A. Förderer et al., A wheat resistosome defines common principles of immune receptor channels. Nature 610, 532–539 (2022).

7. Y.-B. Zhao et al., Pathogen effector AvrSr35 triggers Sr35 resistosome assembly via a direct recognition mechanism. Science Advances 8, eabq5108 (2022).

8. S. Cesari, Multiple strategies for pathogen perception by plant immune receptors. New Phytol 219, 17–24 (2018).

9. J. D. G. Jones, R. E. Vance, J. L. Dangl, Intracellular innate immune surveillance devices in plants and animals. Science 354, aaf6395 (2016).

10. J. Wang et al., Ligand-triggered allosteric ADP release primes a plant NLR complex. Science 364 (2019).

11. J. Wang et al., Reconstitution and structure of a plant NLR resistosome conferring immunity. Science 364 (2019).

12. K. de Guillen et al., Structure Analysis Uncovers a Highly Diverse but Structurally Conserved Effector Family in Phytopathogenic Fungi. PLoS Pathog 11, e1005228 (2015).

13. D. L. Hawksworth, The magnitude of fungal diversity: the 1.5 million species estimate revisited. Mycological Research 105, 1422–1432 (2001).

14. A. R. Bentham et al., A molecular roadmap to the plant immune system. Journal of Biological Chemistry 295, 14916–14935 (2020).

15. D. S. Yu, et al., The structural repertoire of *Aquarium oxysporum* f. sp. *lycopersici* effectors revealed by experimental and computational studies. BioRxiv 10.1101/2021.12.14.472499 (2022).

16. X. Di et al., Structure-function analysis of the *Fusarium oxysporum* Avr2 effector allows uncoupling of its immune-suppressing activity from recognition. New Phytol 216, 897–914 (2017).

17. N. Lazar et al., A new family of structurally conserved fungal effectors displays epistatic interactions with plant resistance proteins. PLoS Pathog 18, e1010664 (2022).

18. S. R. Mark C. Derbyshire, Surface frustration re-patterning underlies the structural landscape and evolvability of fungal orphan candidate effectors. BioRxiv 10.1101/2023.01.06.522876 (2023).

19. M. A. Outram, M. Figueroa, J. Sperschneider, S. J. Williams, P. N. Dodds, Seeing is believing: Exploiting advances in structural biology to understand and engineer plant immunity. Curr Opin Plant Biol 67, 102210 (2022).

20. K. Seong, K. V. Krasileva, Prediction of effector protein structures from fungal phytopathogens enables evolutionary analyses. Nat Microbiol 8, 174–187 (2023).

21. X. Zhang et al., A positive-charged patch and stabilized hydrophobic core are essential for avirulence function of AvrPib in the rice blast fungus. Plant J 96, 133–146 (2018).

22. I. M. Saur et al., Multiple pairs of allelic MLA immune receptor-powdery mildew AVR(A) effectors argue for a direct recognition mechanism. Elife 8 (2019).

23. M. C. Müller, et al., Ancient variation of the *AvrPm17* gene in powdery mildew limits the effectiveness of the introgressed rye *Pm17* resistance gene in wheat. 119, e2108808119 (2022).

24. B. Manser et al., Identification of specificity-defining amino acids of the wheat immune receptor Pm2 and powdery mildew effector AvrPm2. Plant J 106, 993–1007 (2021).

25. X. Lu et al., Allelic barley MLA immune receptors recognize sequence-unrelated avirulence effectors of the powdery mildew pathogen. Proc Natl Acad Sci USA 113, E6486–E6495 (2016).

26. L. Kunz et al., The broad use of the *Pm8* resistance gene in wheat resulted in hypermutation of the *AvrPm8* gene in the powdery mildew pathogen. BMC Biology 21, 29 (2023).

27. S. Bourras et al., Multiple avirulence loci and allele-specific effector recognition control the *Pm3* race-specific resistance of wheat to powdery mildew. Plant Cell 27, 2991–3012 (2015).

28. S. Bourras et al., The AvrPm3-Pm3 effector-NLR interactions control both race-specific resistance and host-specificity of cereal mildews on wheat. Nature Communications 10 (2019).

29. S. Bauer et al., The leucine-rich repeats in allelic barley MLA immune receptors define specificity towards sequence-unrelated powdery mildew avirulence effectors with a predicted common RNase-like fold. Plos Pathogens 17 (2021).

30. C. R. Praz et al., AvrPm2 encodes an RNase-like avirulence effector which is conserved in the two different specialized forms of wheat and rye powdery mildew fungus. New Phytol 213, 1301–1314 (2017).

31. H. G. Pennington et al., The fungal ribonuclease-like effector protein CSEP0064/BEC1054 represses plant immunity and interferes with degradation of host ribosomal RNA. PLoS Pathog 15, e1007620 (2019).

32. L. Frantzeskakis et al., Signatures of host specialization and a recent transposable element burst in the dynamic one-speed genome of the fungal barley powdery mildew pathogen. BMC Genomics 19, 381 (2018).

33. S. Kusch et al., Long-term and rapid evolution in powdery mildew fungi. Molecular Ecology 00 1– 22. (2023).

34. M. C. Müller et al., A chromosome-scale genome assembly reveals a highly dynamic effector repertoire of wheat powdery mildew. New Phytol 221, 2176–2189 (2019).

35. C. Pedersen et al., Structure and evolution of barley powdery mildew effector candidates. BMC Genomics 13, 694 (2012).

36. A. A. Ahmed et al., The Barley Powdery Mildew Candidate Secreted Effector Protein CSEP0105 Inhibits the Chaperone Activity of a Small Heat Shock Protein *Plant Physiology* 168, 321–333 (2015).

37. H. Yuan et al., The powdery mildew effector CSEP0027 interacts with barley catalase to regulate host immunity. Front. Plant Sci. 1967 (2021).

38. W.-J. Zhang et al., Interaction of barley powdery mildew effector candidate CSEP0055 with the defence protein PR17c. Mol. Plant Pathol. 13, 1110–1119 (2012).

39. Z. Li et al., Powdery mildew effectors AVR_A1_ and BEC1016 target the ER J-domain protein *Hv*ERdj3B required for immunity in barley. bioRxiv, 10.1101/2022.04.27.489729 (2022).

40. D. Godfrey et al., Powdery mildew fungal effector candidates share N-terminal Y/F/WxC-motif. BMC Genomics 11, 317 (2010).

41. C. Pedersen et al., Structure and evolution of barley powdery mildew effector candidates. BMC Genomics 13, 694 (2012).

42. L. Holm, P. Rosenstrom, Dali server: conservation mapping in 3D. Nucleic acids research 38, W545–549 (2010).

43. D. Yu et al., TIR domains of plant immune receptors are 2’,3’-cAMP/cGMP synthetases mediating cell death. Cell 185, 2370–2386 e2318 (2022).

44. I. M. L. Saur, S. Bauer, X. Lu, P. Schulze-Lefert, A cell death assay in barley and wheat protoplasts for identification and validation of matching pathogen AVR effector and plant NLR immune receptors. Plant Methods 15, 118 (2019).

45. J. Jumper et al., Highly accurate protein structure prediction with AlphaFold. Nature 596, 583–589 (2021).

46. D. Ortiz et al., The stem rust effector protein AvrSr50 escapes Sr50 recognition by a substitution in a single surface-exposed residue. New Phytol 234, 592–606 (2022).

47. A. R. Bentham, et al., Allelic compatibility in plant immune receptors facilitates engineering of new effector recognition specificities. bioRxiv, 10.1101/2022.10.10.511592 (2022).

48. T. Maekawa et al., Subfamily-Specific Specialization of RGH1/MLA Immune Receptors in Wild Barley. Mol Plant Microbe Interact 32, 107–119 (2019).

49. A. Förderer, D. Yu, E. Li, J. Chai, Resistosomes at the interface of pathogens and plants. Current Opinion in Plant Biology 67, 102212 (2022).

50. P. D. Spanu, Cereal immunity against powdery mildews targets RNase-Like Proteins associated with Haustoria (RALPH) effectors evolved from a common ancestral gene. New Phytol 213, 969–971 (2017).

51. S. He et al., The Secreted Ribonuclease SRE1 Contributes to Setosphaeria turcica Virulence and Activates Plant Immunity. Frontiers in microbiology 13, 941991 (2022).

52. G. J. Kettles et al., Characterization of an antimicrobial and phytotoxic ribonuclease secreted by the fungal wheat pathogen Zymoseptoria tritici. New Phytol 217, 320–331 (2018).

53. B. Yang et al., Fg12 ribonuclease secretion contributes to Fusarium graminearum virulence and induces plant cell death. J Integr Plant Biol 63, 365–377 (2021).

54. W. Kabsch, Xds. Acta Crystallogr D Biol Crystallogr 66, 125–132 (2010).

55. C. Vonrhein et al., Data processing and analysis with the autoPROC toolbox. Acta Crystallographica Section D 67, 293–302 (2011).

56. P. S. Bond, K. D. Cowtan, ModelCraft: an advanced automated model-building pipeline using Buccaneer. Acta Crystallographica Section D 78, 1090–1098 (2022).

57. P. Emsley, B. Lohkamp, W. G. Scott, K. Cowtan, Features and development of Coot. Acta Crystallogr D Biol Crystallogr 66, 486–501 (2010).

58. D. Liebschner et al., Macromolecular structure determination using X-rays, neutrons and electrons: recent developments in Phenix. Acta Crystallographica Section D 75, 861–877 (2019).

59. E. F. Pettersen et al., UCSF ChimeraX: Structure visualization for researchers, educators, and developers. Protein Sci., 30, 70–82 (2021).

60. X. Robert, P. Gouet, Deciphering key features in protein structures with the new ENDscript server. Nucleic acids research 42, W320–324 (2014).

61. A. Himmelbach et al., A set of modular binary vectors for transformation of cereals. Plant Physiol 145, 1192–1200 (2007).

62. T. Nakagawa et al., Improved Gateway binary vectors: high-performance vectors for creation of fusion constructs in transgenic analysis of plants. Biosci Biotechnol Biochem 71, 2095–2100 (2007).

63. A. V. Garcia et al., Balanced nuclear and cytoplasmic activities of EDS1 are required for a complete plant innate immune response. PLoS Pathog 6, e1000970 (2010).

64. K. Norkunas, R. Harding, J. Dale, B. Dugdale, Improving agroinfiltration-based transient gene expression in Nicotiana benthamiana. Plant Methods 14, 71 (2018).

65. M. Cianci et al., P13, the EMBL macromolecular crystallography beamline at the low-emittance PETRA III ring for high- and low-energy phasing with variable beam focusing. Journal of Synchrotron Radiation 24, 323–332 (2017).

66. A. A. McCarthy et al., ID30B -a versatile beamline for macromolecular crystallography experiments at the ESRF. Journal of Synchrotron Radiation 25, 1249–1260 (2018).

